# Chromatin and gene-regulatory dynamics of human pulmogenesis by single cell multiomic sequencing

**DOI:** 10.1101/2025.09.24.678382

**Authors:** Samuel H. Kim, Soon Il Higashino, Lea C. Steffes, Li Li, Anca M. Pasca, Maya Kumar, Christin S. Kuo, Mark A. Krasnow, William J. Greenleaf

## Abstract

Human lung development is governed by complex gene regulatory networks that orchestrate cellular differentiation and organogenesis. We present a single cell multiomic atlas of human pulmogenesis, simultaneously capturing both the chromatin accessibility profile and the transcriptome from each cell across fetal lungs spanning from post-conception weeks (PCW) 12 to 23. We identified 44 distinct developing cell clusters and mapped 581,745 candidate cis-regulatory elements and nominated 121,486 non-redundant peak-to-gene linkages. We identify highly regulated genes (HRGs) and the cognate highly regulating peaks (HRPs) that describe the most salient regulatory gene programs and developmental enhancer sites for each cell type. Trajectory analysis along with interpretable cell type specific convolutional neural network models were developed to delineate dynamic regulatory programs driving key developmental transitions, including aerocyte and arterial differentiation and alveolar formation. Furthermore, we identified distinct vascular smooth muscle subpopulations with unique spatial associations to either arterial or venous structures with reciprocal signaling within each niche. We also uncovered the regulatory modules of surfactant production in alveolar progenitors, implicating a direct role for the glucocorticoid receptor alongside novel transcription factors. Finally, using cell type specific models linking DNA sequence to chromatin accessibility we prioritize variants associated with impaired pulmonary function or disease and nominate mechanisms of motif disruption. Overall, our multiomic atlas deepens our understanding of the gene-regulatory architecture underlying human lung development and provides a valuable resource for the community to dissect the cellular and molecular programs of pulmonary physiology and disease at the cellular and nucleotide precision.

## Introduction

Pulmogenesis begins with ventral foregut endodermal outpouching and branching morphogenesis to form the conducting airways of the lungs (Metzger et al., 2008; Zepp and Morrisey, 2019). At the cellular level, highly specialized epithelial and endothelial cell types supported by stromal structures develop to form alveoli, with their air-blood interfaces specialized for postnatal gas exchange (Gillich et al., 2020; Kumar et al., 2014; Travaglini et al., 2020). Recent efforts have deeply characterized the developmental gene expression profiles of some of these phenotypically diverse cell types using single cell RNA sequencing (He et al., 2022; Quach et al., 2024; Sountoulidis et al., 2023). However, there yet remains a critical knowledge gap in the molecular drivers and regulatory programs of the developing cell types. Cell type specific expression and activity of transcription factors (TFs) on accessible DNA elements compose the regulatory programs that direct differentiation of diverse cell types from the blueprint of an identical genome (Klemm et al., 2019). This knowledge gap in understanding and defining the drivers and regulatory programs of pulmogenesis might be bridged by identifying transcription factor expression and activity for each cell type, connecting DNA regulatory element accessibility to cognate target gene expression, and specifying TF motifs within those DNA regulatory elements to deduce the interaction of TFs and regulatory elements, and provide a comprehensive framework of the master regulators and gene expression programs of each lung cell type.

Single cell methods for probing chromatin accessibility, such as the assay for transposase-accessible chromatin (scATAC-seq), provide a cell type-resolved, genome wide profile of accessible DNA, which can be used to infer the chromatin activity of TF binding motifs during differentiation (Buenrostro et al., 2015). Methods for simultaneously capturing both the chromatin accessibility and the transcriptome from the same cell have enabled the identification of DNA elements with chromatin accessibility correlated with gene expression of nearby genes, especially along continuous trajectories in development (Kartha et al., 2021; Ma et al., 2020). Neural network models trained on these cell type specific accessibility profiles have recently enabled motif predictions in regulatory DNA elements by inferring base pair resolution contribution to accessibility within each DNA element (Avsec et al., 2021a; Pampari et al., 2025). These predictions have further allowed for the prioritization of disease implicated sequence variants and nomination of motifs disrupted by the mutation (Ameen et al., 2022; Corces et al., 2020; Ober-Reynolds et al., 2023; Trevino et al., 2021).

To map the gene regulatory programs in pulmogenesis, we generated a single cell multiomic atlas by simultaneously capturing chromatin accessibility and RNA expression from the same cell across human fetal lung samples from post conception week (PCW) 12 to 23. We identified putative enhancers with chromatin accessibility that is strongly correlated with target gene expression and nominated “highly regulated genes” (HRGs) for each cell type. We trained and interpreted convolutional neural network models from the cell type specific chromatin accessibility profiles to identify TF motifs that are driving chromatin accessibility in regulatory elements. We then applied these insights across each tissue compartment of the lung. In the endothelial compartment, we identified developmental trajectories for aerocytes and arterial cells and nominated previously disease implicated and novel TF regulators with motif accessibility and expression correlated along the trajectories. In the stromal compartment, we identified anatomically, functionally, and developmentally distinct novel subtypes of vascular smooth muscle cells. In the epithelial compartment, we nominated regulators of surfactant production at the early stages of alveologenesis, implicating a direct role for the glucocorticoid receptor and additional transcription factors with therapeutic implications in lung maturation in premature births. Lastly, we used a sequence to accessibility classifier trained on the chromatin accessibility profiles to prioritize variants of pulmonary function and disease in a cell type specific manner and used this interpretation engine to dissect the potential pathophysiological mechanisms of asthma variants.

## Results

### Single cell multiomic atlas of human pulmogenesis

We used the 10x Genomics Multiome platform to simultaneously assay both the transcriptome and the chromatin accessibility profile from cells derived from seven primary lung samples from PCW 12, 14, 15, 16, 19, 21, and 23 (Figure 1A, C). Additional annotations for anatomic location (proximal, distal, and lobe identity) were collected when available. After quality control and filtering (Figure S1), we obtained 69,330 cells with both high-quality transcriptome and the chromatin accessibility data (Figure 1B).

**Figure 1:**
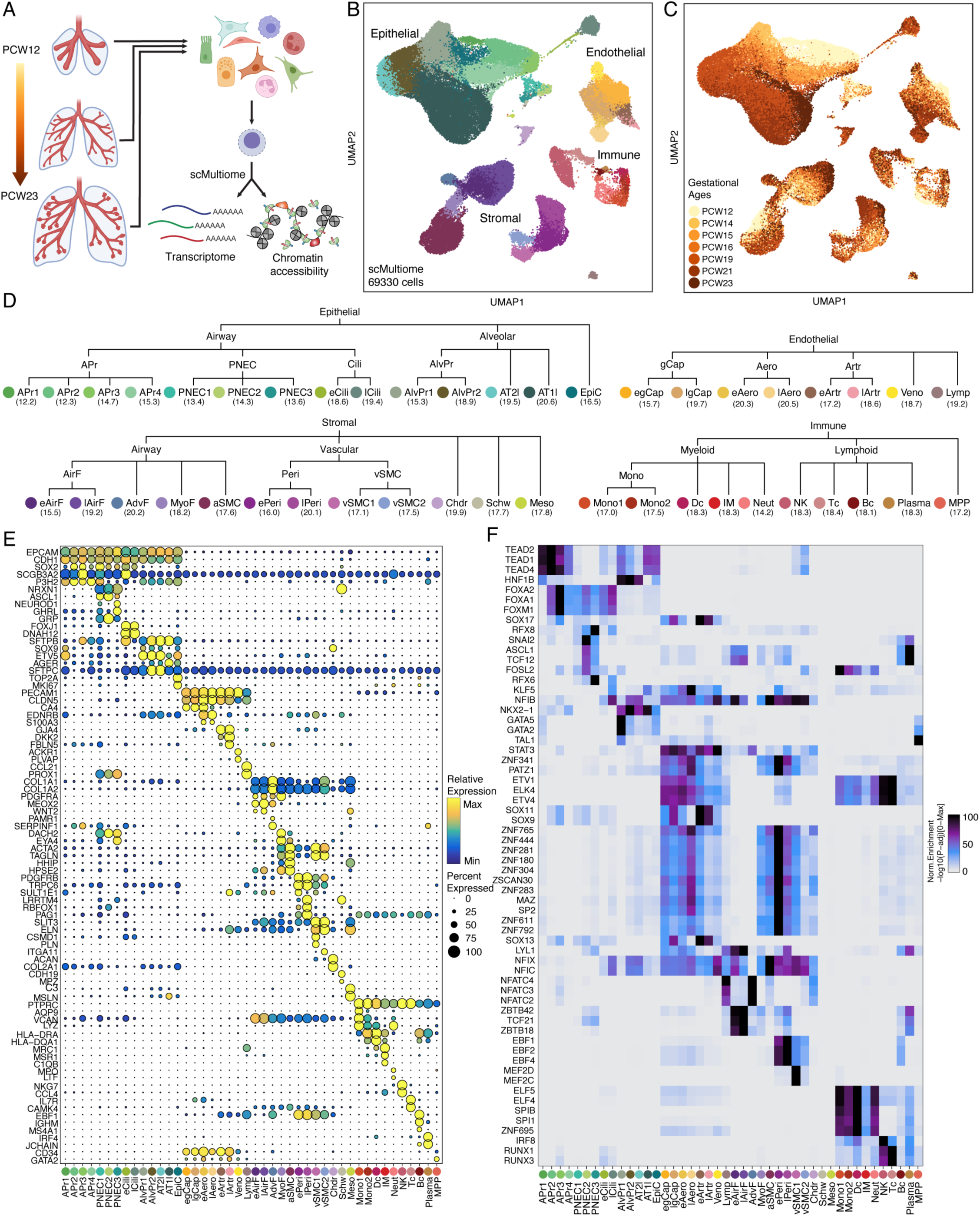
Overview of the single cell multiomic atlas of human pulmogenesis. A) Schematic of sample collection and single cell multiomic profiling. B) UMAP representation of cells colored based on cluster identity. C) UMAP representation of cells colored based on gestational age by post-conception week (PCW). D) Hierarchical organization of cluster assignments for each compartment with mean gestational age for each cluster. E) Marker gene expression for each cluster. Color of the dot indicates the relative expression of the gene across clusters. Dot size indicates percent of cells within each cluster expressing the gene. F) Hypergeometric enrichment of motifs of expressed transcription factors in each cluster.

We annotated cell types of the developing human lung with an iterative subclustering strategy (Figure S1A). After the initial consensus variable gene selection, dimensionality reduction, and clustering at the level of the whole lung, we separated clusters into compartments based on compartment-specific marker gene expression (Epithelial: EPCAM, CDH1; Endothelial: PECAM1, CLDN5; Stromal: COL1A1, COL1A2, ACTA2, TAGLN; Immune: PTPRC/CD45) (Travaglini et al., 2020). Instead of reference integration that can result in spuriously granular annotations without biological significance, we focused on defining cell types based on known markers that were validated in literature or novel markers related to a *bone fide* biological function. In total, 44 cell types of the developing human lung were annotated at high resolution across four compartments (Figure 1D). Each cluster included cells across multiple samples and time points (Figure S2). We identified developmental intermediate cell states based on known markers, including early aerocytes (eAero) that express both CA4 and EDNRB (Gillich et al., 2020). We also identified novel marker genes associated with later stages of pericyte (lPeri) differentiation (LRRTM4, RBFOX1, and PAG1).

From the corresponding chromatin accessibility profiles of each cluster, we identified 581,745 peaks of open chromatin as candidate cis-regulatory elements (CREs). Marker peaks were identified for each cluster and transcription factor motif enrichment in cluster specific marker peaks were calculated. (FDR ≤ 0.01 and Log2FC ≥ 1) (Figure 1F). Motifs of transcription factors previously described as necessary for lung epithelial differentiation (NKX2-1, FOXA2) were enriched in marker peaks for alveolar and airway progenitors (Minoo et al., 1999; Wan et al., 2004). Motifs of transcription factors described in other organs as regulating differentiation of vascular smooth muscle cells and pericytes, such as MEF2C and EBF1, were also enriched in the respective marker peaks for pulmonary smooth muscle cells and pericytes (Materna et al., 2018; Pagani et al., 2021).

We found that multiome data provides more granular cell type specific peak calling for chromatin accessibility compared to using just scATAC-seq, because the cell type identity can be assigned directly using the gene expression information without relying on promoter and gene body accessibility as a proxy for gene expression. To illustrate the utility afforded by multiomic data, we treated our multiome data as two singleome datasets, assigned cluster identities based on chromatin-accessibility-based gene scores derived from promoter accessibility, then reintegrated the data with canonical correlation analysis (Granja et al., 2021; Stuart et al., 2019). Calculating the Jaccard similarity index based on the ground truth assignments of the cell barcodes showed that multiple distinct cell type identities across all compartments (i.e. AT1l vs. AT2l, all endothelial cell types) were lost when integration was performed across the two singleomes (Figure S3A). Furthermore, cell states along continuous trajectories (i.e. eAero and lAero) were not observed. We then assessed whether incorporating back the true multiomic data by assigning cluster identities directly using gene expression would provide more granular chromatin accessibility profiles that capture cell type specific accessibility. For this, we performed the same integration but with gene scores calculated from chromatin accessibility profiles derived from multiomic assignment of clusters. By treating the data as true multiome, the chromatin accessibility profiles of each cluster recapitulated almost all the cluster identities derived purely from gene expression (Figure S3A). This analysis demonstrates how multiomic data allows more fine-grained cluster calling with the RNA modality, which then allows for more fine-grained analysis of the linked ATAC-seq data.

### Linking gene expression regulation to chromatin accessibility across human pulmogenesis

To understand the transcriptional regulation that drives cell type specification in human lung development, we identified chromatin accessibility peaks that were correlated to expression of nearby genes. A total of 121,486 non-redundant peak-to-gene linkages were detected between 80,121 accessible CREs and 13,260 expressed genes (Figure 2A). K-means clustering based on accessibility at the linked CREs revealed the heterogeneity of the CRE usage for cell type and compartment specific gene expression. The CREs linked to nearby genes represent 13.7% of all peaks. While developmental enhancer validation experiments at cell type resolution are limited, we compared our nominated linked peaks against specific enhancers validated in developing zebrafish. Hu et al. tested all conserved noncoding regions 100kb upstream of the ACAN locus, a structural protein for cartilage, via a reporter assay throughout zebrafish development. We observed good overlap between linked peaks identified for ACAN locus and these developmentally validated enhancer sequences (Figure 2B) (Hu et al., 2012). Furthermore, enhancers from Hu et al. that were only transiently active during development and became inactive later in development were also identified from peak to gene linkage analysis, consistent with the developmental context of the chondrocytes in this atlas.

**Figure 2:**
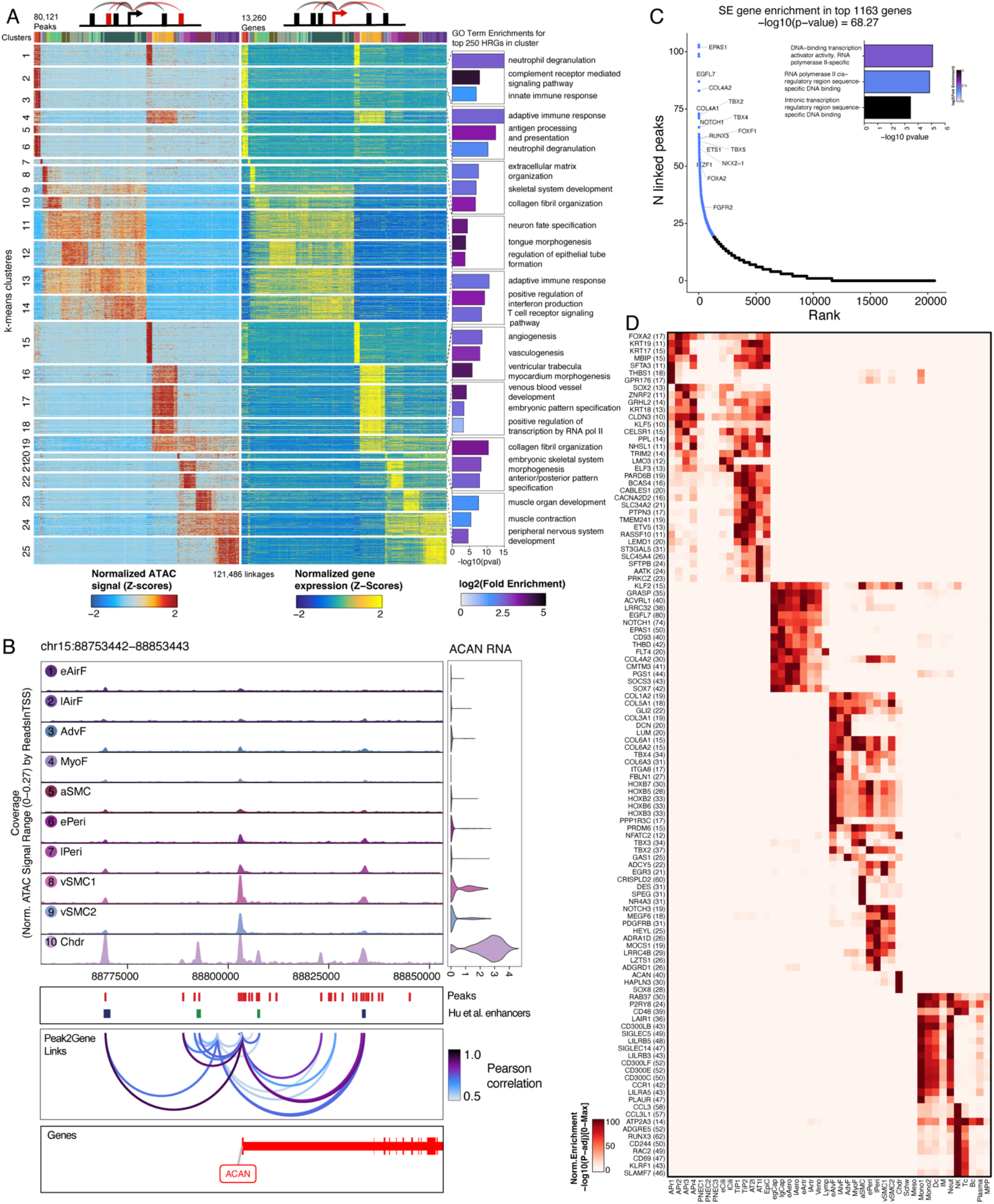
Multiomic gene regulatory dynamics in human lung. A) Heatmap of 121,486 significant peak-to-gene linkages between 80,121 accessible candidate cis-regulatory elements (cCRE) and 13,260 expressed genes. Each row in the paired heatmap represents one cCRE and its linked gene. For visualization, the rows were clustered using k-means clustering (k = 25). Top 3 enriched gene ontology terms for top 250 highly regulated genes (HRGs) are shown for selected clusters. B) Peak2gene linkages nominate cCREs for chondrocyte specific ACAN expression. Enhancer regions confirmed in zebrafish for cartilage expression from Hu et al. (2013) highlighted in dark blue. Enhancer regions only active during development are highlighted in dark green. Violin plot of ACAN RNA counts are shown for each cluster on the right panel. C) Ranked list of highly regulated genes (HRGs). Each gene is ranked based on the number of peak2gene linkages. 1163 HRGs have more than 20 peak2gene linkages. Top 3 enriched gene ontology terms for top 100 HRGs are shown in the inset. D) Heatmap of enrichment of highly regulating peaks (peaks that are linked to HRGs) in marker peaks for a given cluster against the union set of all marker peaks.

Next, we identified set of highly regulated genes (HRGs) with many CREs with accessibility correlated to its expression (Figure 2C), representing genes with the most regulation by CREs. We identified 1163 HRGs based on genes that have at least 20 peaks linked to its expression. These HRGs include key transcription factors necessary for lineage specification (NKX2-1, FOXA2, RUNX3) as well as genes involved in signal transduction for specific cellular functions (NOTCH1, FGFR2). GO term enrichment for the top HRGs in each k-means cluster highlights terms relevant for cell type function, such as “adaptive immune response” for T cells and “collagen fibril organization” for fibroblasts (Figure 2A). Top GO term enrichments for the top 100 HRGs across all cell types are related to transcription factor binding and gene regulation. The HRGs reveal the most salient genes for cell type specification and function.

Then, we identified set of highly regulating peaks (HRPs) that are linked to the expression of HRGs. To identify which cell types are accessible at the HRPs, we calculated enrichment of HRPs in each cluster’s marker peak against the union set of all marker peaks. This strategy revealed linkages to both cell type specific HRGs (ACAN in chondrocytes, DES in airway smooth muscle cells) but also modules of HRGs common to a given lineage of multiple cell types (FOXA2 across epithelial progenitors, KLF2 across endothelial progenitors) (Figure 2D). Together, we mapped out the gene regulatory programs that underly pulmogenesis by identifying HRGs and HRP usage for each cluster and lineage.

### Regulators of pulmonary endothelial differentiation

Pulmonary endothelium, particularly at the air-blood interface, is highly specialized to maximize surface area and minimize diffusion distance to enable efficient gas exchange (Gillich et al., 2020). To dissect the regulatory programs defining these endothelial cell types, we first subclustered the endothelial compartment (Figure 3A), resulting in 8 subclusters. To identify the common progenitor within the endothelial compartment, we performed differential abundance testing on the subclustered dataset to identify clusters that were over-represented in earlier time points. egCaps showed significant enrichment for earlier time points as well as eArtr cells (Figure S5A). egCaps lacked any of the terminal endothelial cell type markers and expressed CA4 (Figure 1E). To define a trajectory of endothelial differentiation, we applied the optimal transport algorithm implemented within CellRank (Lange et al., 2022; Schiebinger et al., 2019a). egCaps were identified as the initial state with lAero, lArtr, Veno, and lgCap as the 4 terminal states (Figure S5B). Fate probabilities from egCaps to the four terminal states (Figure 3B) suggest that egCaps represent the capillary plexus cells previously described to be early endothelial capillaries that cover the developing airway and are capable of further differentiating into the vascular cell types of the lung (Gillich et al., 2020).

**Figure 3:**
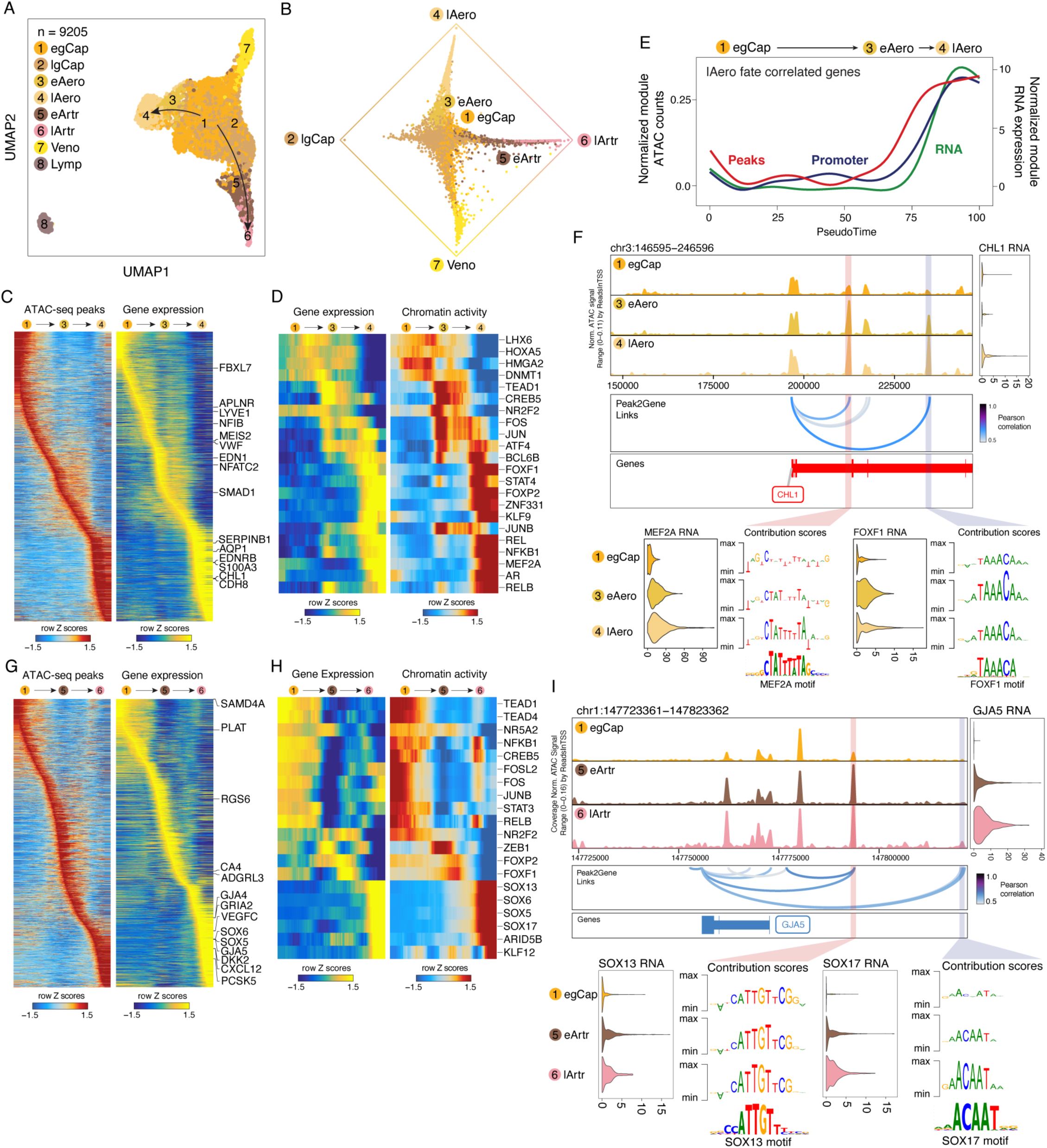
Dissecting regulators of endothelial differentiation in human pulmogenesis. A) UMAP representation of cells in the endothelial compartment. B) Circular projection (Lange et al., 2022; Velten et al., 2017) of endothelial cells according to fate probabilities towards terminal macrostates, colored according to cluster identity. C) Heatmap of accessible peaks and gene expression along aerocyte differentiation trajectory. Marker genes are labeled accordingly. D) Paired heatmaps of transcription factors along the aerocyte differentiation trajectory whose expression is correlated with its motif activity. E) Sequential peak accessibility (red), promoter accessibility (blue), and gene expression (green) along pseudotime for aerocyte trajectory. F) ChromBPNet predicted transcription factor motif (MEF2A, FOXF1) accessibility in peaks linked to expression of a gene correlated with aerocyte trajectory (CHL1). G) Heatmap of accessible peaks and gene expression along arterial differentiation trajectory. Marker genes are labeled accordingly. H) Paired heatmaps of transcription factors along the arterial differentiation trajectory whose expression is correlated with its motif activity. I) ChromBPNet predicted transcription factor motif (SOX13, SOX17) accessibility in peaks linked to expression of a gene correlated with arterial trajectory (GJA5).

Next, we sought to identify regulators along the trajectories from egCaps to aerocyte or arterial states. First, we identified trajectories across cells by aligning the sequence of clusters based on fate probabilities (Lange et al., 2022; Weiler et al., 2024). Leveraging the multiomic data, we identified a set of variable peaks and genes across the trajectories, which highlighted previously described key marker genes for aerocytes (EDNRB, S100A3) (Gillich et al., 2020) and arterial cell types (GJA5, DKK2) (Travaglini et al., 2020) (Figures 3C, G) as well as nominating new marker genes. To identify regulators along this trajectory, we correlated transcription factor expression to its motif accessibility to nominate candidate transcription factors regulating this differentiation trajectory (Figure 3D, H). Along the aerocyte trajectory, we identified FOXF1 as well as other candidate transcription factors such as MEF2A. FOXF1 mutations have been implicated in alveolar capillary dysplasia (Sen et al., 2013), which leads to improper pulmonary capillary distribution and structure during development. Along the arterial trajectory, multiple SOX transcription factors were nominated, such as SOX17, which has previously been shown to be necessary for arterial differentiation (Corada et al., 2013). Next, we assessed the temporal sequence of regulatory events establishing the aerocyte state. Along the pseudotime of the aerocyte trajectory, we noted that the distal CREs linked to aerocyte genes gained accessibility first, followed by increased promoter accessibility, followed by increased expression of linked genes (Figure 3E).

To further dissect the activity of transcription factors at CREs, we trained ChromBPNet convolutional neural networks to learn cluster specific Tn5 bias-factorized mapping from the DNA sequences of accessible peaks to a corresponding base-resolution chromatin accessibility profile (Pampari et al., 2025). In model performance testing across 5-fold chromosome hold-out cross-validation scheme, we observed high Spearman correlation (Figure S4A) between observed and predicted counts in peaks and low median Jenson-Shannon distance (Figure S4B) between observed and predicted base resolution profiles. We next interpreted each cluster-specific model to calculate the count and profile contribution scores at each base pair of accessible peaks using DeepLIFT (Avsec et al., 2021b; Shrikumar et al., 2017). Interpretation of these models identified short sequences with high contribution scores that resemble known TF binding motifs and likely representing transcription factor binding activity at these sites (Ameen et al., 2022).

Because experimental dissection of specific TF binding within enhancers in primary developmental cell types is limited, we compared our model predictions to a conserved enhancer of PROX1 where specific TF binding sites have been characterized and confirmed with cell type resolution in embryonic mice (Kazenwadel et al., 2023). Using our approach, we observed increased contribution scores in lymphatic cells at the exact NFATC1 motif previously shown to bind NFATC1 and function as a validated conserved enhancer for PROX1 regulation in lymphangiogenesis (Figure S5C). Additionally, we observed within the developing human lung that PROX1 expression was present in pulmonary neuroendocrine cells (PNECs) as previously reported in mouse pulmonary neuroendocrine cells and in carcinoid cells (McGovern et al., 2010; Sakurai et al., 2024). The NFATC1 motif, active in lymphatic cells, was not predicted to be contributing accessibility at the same enhancer in PNECs but at a nearby HNF1B motif. These observations suggest that cell type specific enhancer usage can occur through multiple TF binding sites within the same enhancer.

To infer which transcription factors are active near terminal state correlated genes of endothelial cell types, we used the chromBPNet model predictions in linked CREs. First, we identified CHL1 as a gene highly correlated in expression along the aerocyte trajectory and identified two linked peaks near the CHL1 gene that showed increase in accessibility along development (Figure 3F). In these CREs, we predict concurrent increase in accessibility for MEF2A and FOXF1 motifs, two of the candidate transcription factors nominated by its correlated expression and motif activity (Figure 3F). As expected, expression of MEF2A and FOXF1 increases along the differentiation trajectory. Along the arterial differentiation, we identified two linked peaks to GJA5, a marker gene for arterial cells with expression correlated with the arterial trajectory. These CREs gain accessibility along arterial development (Figure 3I). Here the SOX13 and SOX17 motifs are predicted to gain accessibility along development with concordant increase in expression. This map of predicted transcription factor motifs along with peak to gene linkages provide a systematic approach to nominate candidate transcription factors and their targets genes and sites of regulation.

### Novel vascular smooth muscle cell types in human pulmogenesis

The stromal compartment represents the second most abundant compartment in the developing human lung and provides diverse structural, contractile, and signaling functions to guide development and later enable respiration (Kumar et al., 2014). After subclustering cells within the stromal compartment, we identified two distinct vascular smooth muscle clusters (Figure 4A) that we name vSMC1 and vSMC2. Both vSMC1 and vSMC2 express contractile genes (ACTA2, TAGLN) and PDGFRB (Figure 1E), and the ratio of vSMC1 to vSMC2 cells does not change throughout the developmental window we examined (Figure S6B). GO term enrichment for differentially expressed genes showed that vSMC1-specific genes are enriched for contractile functions while vSMC2-specific genes are enriched for extracellular matrix organization (Figure 4B). Using the chromatin accessibility profiles for the two distinct vSMCs, we identified differentially expressed transcription factors with correlated motif activity and expression (Figure 4C-D). For vSMC1, MEF family transcription factors (specifically MEF2C and MEF2B) were both specific to vSMC1 and its expression and motif activity correlated for this cell type. MEF family transcription factors have been well characterized in cardiac development in regulating genes for establishing contractile function (Desjardins and Naya, 2016; Materna et al., 2018). In contrast, for vSMC2, GATA6 was differentially expressed with motif activity correlated with its expression. While previously implicated in maintenance of a terminal smooth muscle state (Rzucidlo et al., 2007), a clear role for GATA6 in developing smooth muscle cells is yet to be established.

**Figure 4:**
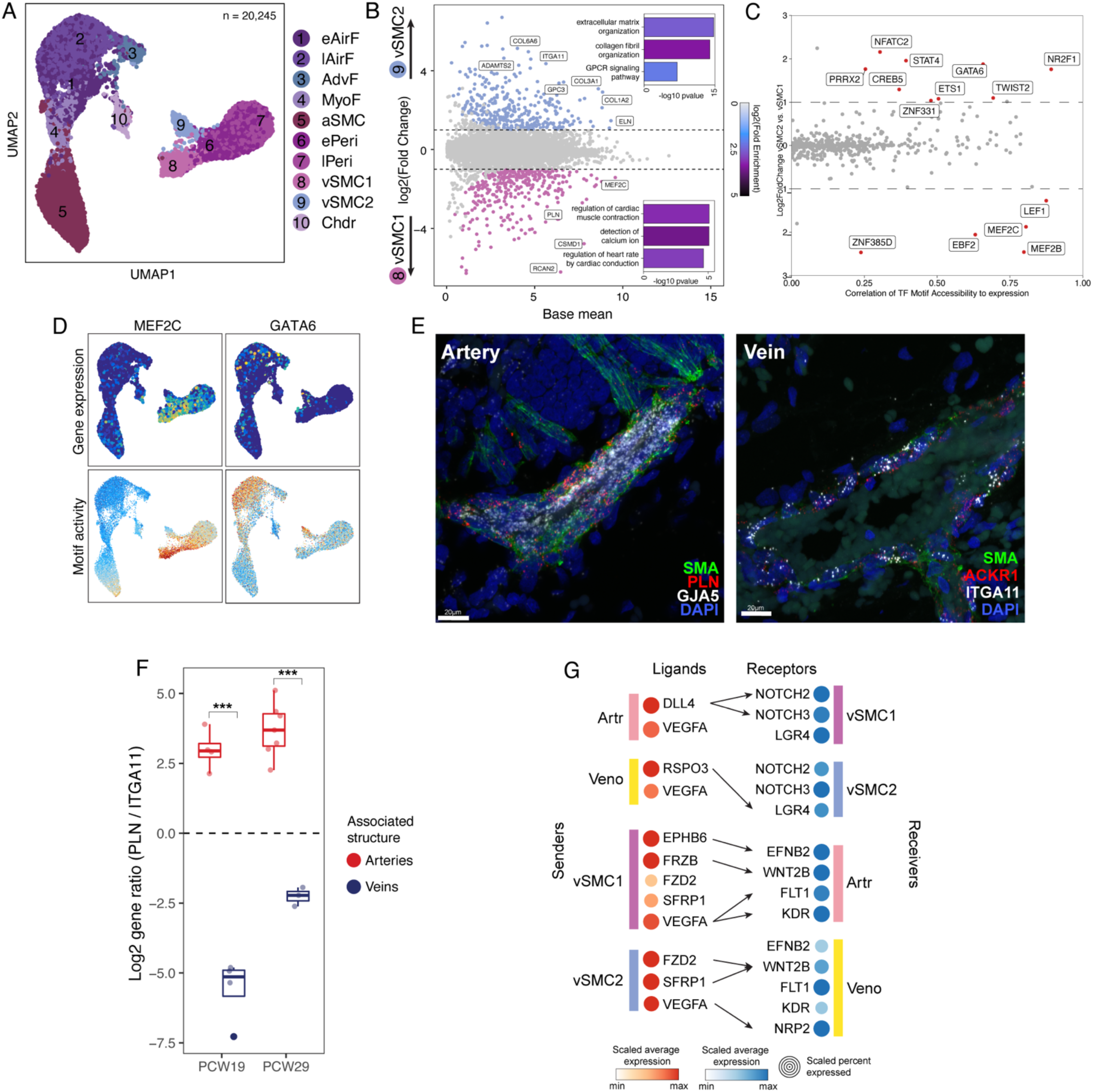
Novel vascular smooth muscle cell types in pulmogenesis. A) UMAP representation of cells in the stromal compartment. B) Differential gene expression between vSMC1 vs. vSMC2. GO term enrichment for differentially expressed genes are shown in the inset bar plot. C) Nominating transcription factor usage between vSMC1 and vSMC2. On x-axis, each transcription factor is plotted based on correlation of its motif accessibility to expression. On y-axis, log 2-fold change for expression of that transcription factor in vSMC1 vs. vSMC2. Transcription factors are colored red and labeled if they are specific to a given cluster in expression (log2FC > 1 or < -1) and correlation of its motif accessibility to expression is > 0.2. D) UMAP representation with cells colored based on expression and motif activity of the transcription factors (MEF2C, GATA 6). E) Representative IF-FISH images of PCW 19 artery and vein with surrounding vascular smooth muscle cells. Smooth muscle actin (SMA) antibodies are pseudocolored to green, FISH probes for ACKR1 (Vein) or PLN (vSMC1) in red, GJA5 (Arterial) or ITGA11 (vSMC2) in white, and DAPI in blue. F) Quantification of PLN to ITGA11 expression ratio in vascular smooth muscle cells at PCW19 and PCW29. Associated structures were determined based on morphology and expression of GJA5 for arteries and ACKR1 for veins. Welch’s two sample t-test was performed for each PCW, p-value for PCW19: 2.55x10^-5^, PCW29: 7.27x10^-3^. (*** p < 0.01). G) Ligand-receptor co-expression analysis using CellphoneDB v3 between vascular smooth muscle cells and endothelial cell types. Dot size depicts scaled percent expression and color depicts scaled average expression of the ligand or the receptor. Arrows are shown for statistically significant (p < 0.05) ligand-receptor coexpression compared to a null distribution of co-expression observed from randomly permuted cluster labels.

To examine the spatial distribution of the two vSMC cell types, we performed IF-FISH with marker genes identified from differential gene expression analysis (Figure 4E). We observed a clear localization of PLN positive cells (a marker for vSMC1 cells) at arteries compared to ITGA11 positive cells (a marker for vSMC2 cells) surrounding venous structures at two distinct developmental timepoints (Figure 4F). Co-staining of vSMC markers with endothelial markers confirmed the identity of the endothelial structures (Figure 4E, S6D). Thus, vSMC1s preferentially surround arterial endothelium while vSMC2s preferentially surround venous endothelium.

Given this highly specific proximity between vSMC1 to arterial endothelium and vSMC2 to venous endothelium, we hypothesized that reciprocal signaling within each niche may be driving the respective cell type specification. Therefore, we performed ligand-receptor analysis to nominate signaling programs specific to these niches. We identified multiple signaling co-expression of ligands and receptors between cognate endothelial and vSMC pairs through NOTCH, WNT, and VEGF signaling pathways (Figure 4G). We found that NOTCH2 and NOTCH3 are expressed in both vSMCs, however, the DLL4 ligand is only expressed in arterial endothelial cells. Of note, NOTCH3 and NOTCH2 signaling have been shown to be necessary for vascular smooth muscle development in mouse knockout models, specifically for arterial endothelium (Domenga et al., 2004; Wang et al., 2012). Furthermore, DLL4 interaction with NOTCH3 has been shown to be necessary in vasculogenesis around tumors in murine sarcoma models (Stewart et al., 2011). Therefore, the restrictive expression of DLL4 to arterial endothelial cells may promote the vSMC1 state compared to the vSMC2 state. We also found both vSMC1 and vSMC2 cells express VEGFA but arterial endothelial cells express more of the highly active receptor KDR (VEGFR2). VEGFA-KDR signaling has been shown to be essential for angiogenesis in mouse knockout and mutant models (Sakurai et al., 2005; Shalaby et al., 1995) and VEGFA is upstream of DLL4 expression (Wythe et al., 2013), suggesting that the reciprocal signaling between arterial endothelial cells and vSMC1 may be further reinforcing the differences between artery and vein endothelium. This analysis suggests that reciprocal signaling between the specific pairings of vSMCs and endothelial cells are restricted by the spatial niches the cell types occupy.

Overall, the results demonstrate two distinct subpopulations of vSMCs in the developing human lung, with distinct locations, signaling, cellular functions, and transcriptional regulatory programs. We propose the name arterial vascular smooth muscle cell (vSMC-a) for vSMC1s with more contractile programs and located near arterial endothelium and the name venous vascular smooth muscle cell (vSMC-v) for vSMC2s with more structural programs and located near venous endothelium.

### Regulators of surfactant production in human pulmogenesis

Proper surfactant production is necessary for postnatal respiration with complications accounting for significant mortality and morbidity in premature births(Berger et al., 2024; Gruenwald, 1947; Holme and Chetcuti, 2012). We aimed to interrogate our open chromatin data set along alveolar progenitor trajectory to identify regulators of surfactant production. First, we identified 14 cell clusters in the epithelial compartment (Figure 5A). Potential alveolar progenitor (AlvPr) populations were identified based on high SOX9 expression without SOX2 expression and low level of SFTPB expression (Danopoulos et al., 2017; Rockich et al., 2013). We identified alveolar differentiation trajectories from AlvPr1 to AT2l and AT1l clusters as previously described for endothelial differentiation trajectories (Figure 5B). To identify the genes most relevant for lamellar body function and earliest stages of surfactant production, we examined the expression of genes from GO terms for surfactant production (GO:0043129, GO:0042599, GO:0097208, GO:0097233) along the developmental trajectory towards AT2l cluster (Figure 5C). Surfactant genes (SFTPA1, SFTPA2, SFTPB, SFTPC, SFTPD) as well as genes essential for lamellar body function (ABCA3, LAMP3, and CTSH) increase in expression along this trajectory. To identify the enhancers regulating these genes, accessibility in peaks linked to expression of these genes were summed and normalized as a peak module score (Figure 5D). Total accessibility in peaks linked to surfactant genes and lamellar body genes increases along the clusters developing AT2l cells. Motifs enriched in these linked peaks (Figure 5E) include NR3C1, the glucocorticoid receptor. Given the clinical practice of betamethasone treatment in preterm labor to accelerate surfactant production in premature infants, these findings provide a potential direct mechanistic explanation underlying this now decades-long life-saving practice. Furthermore, we nominate additional transcription factors, such as SOX4, SOX11, and EHF as well as the mineralocorticoid receptor (NR3C2) as potential regulators of the earliest stages of surfactant production and lamellar body function.

**Figure 5:**
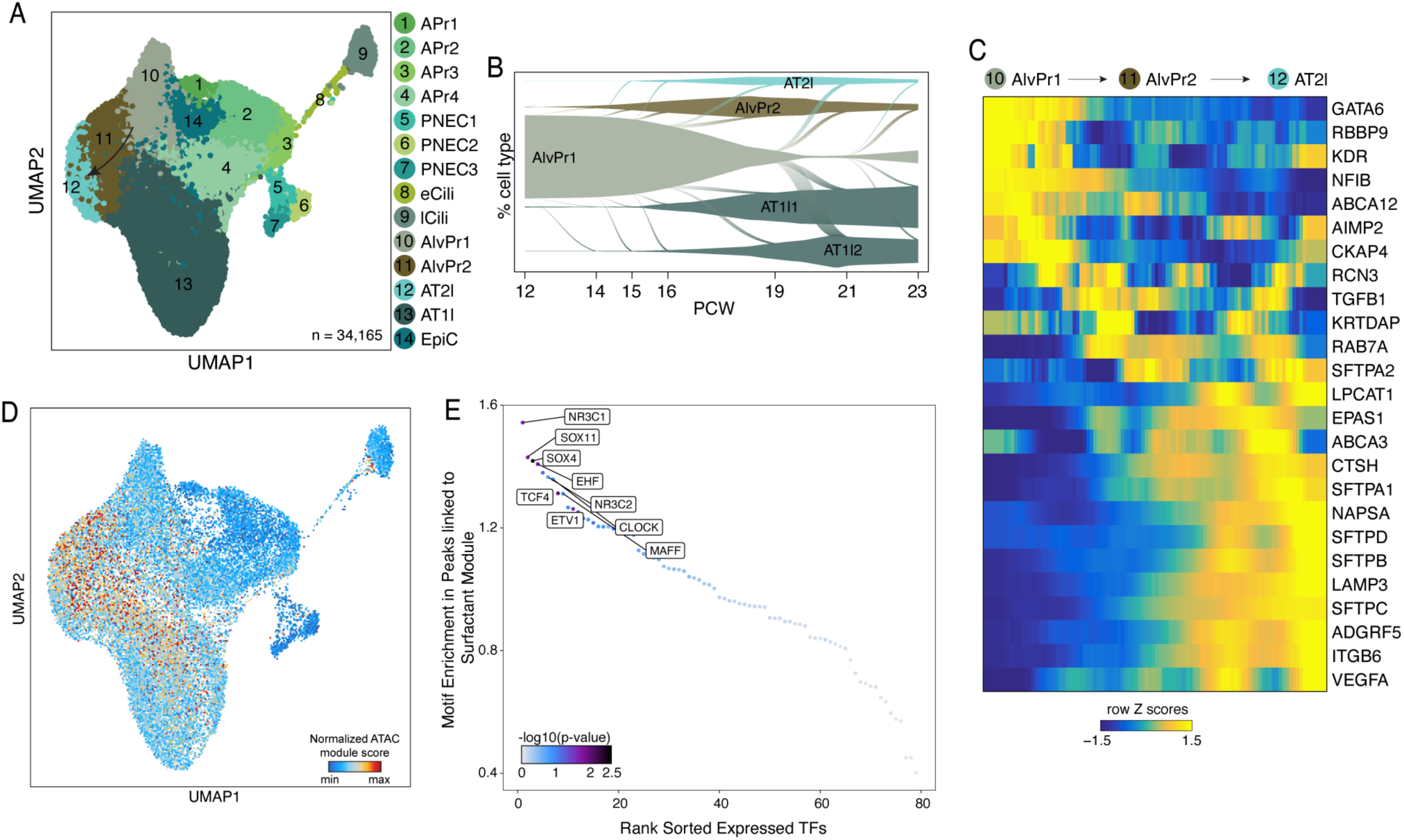
Regulators of surfactant production along epithelial differentiation trajectory. A) UMAP representation of epithelial cells colored based on cluster identity. B) Vein plot (Mittnenzweig et al., 2021) describing flow based on optimal transport maps from AlvPr1 to AT1 vs. AT2. C) Expression of genes from GO terms for surfactant production (GO:0043129, GO:0042599, GO:0097208, GO:0097233) along the differentiation trajectory to AT2l. D) UMAP representation with each cell colored based on accessibility in peaks linked to surfactant module gene expression. E) Motif enrichment of expressed transcription factors within peaks linked to surfactant module genes.

### Nominating candidate causal variants in pulmonary disease

Given that disease-associated single nucleotide polymorphisms (SNPs) are enriched in non-coding enhancer elements, (ENCODE Project Consortium, 2012) we sought to nominate candidate causal variants in pulmonary traits with our multiomic atlas of pulmogenesis. First, we performed linkage disequilibrium score regression (Bulik-Sullivan et al., 2015) using GWAS summary statistics across pulmonary and non-pulmonary traits and examined that cell type-specific peaks were enriched in variants associated with these traits (Figure 6A). Variants associated with non-pulmonary traits were not enriched across the cluster specific peaks in human lung development, as expected. In contrast, variants associated with lung function such as forced vital capacity (FVC) of the lung and the ratio of forced expiratory volume in first second to FVC (FEV1/FVC) were significantly enriched in cell type specific peaks across the stromal clusters as well as epithelial clusters involved in alveolar function (AT1l and AT2l). Asthma variants, including childhood onset asthma, were enriched in peaks for immune clusters, specifically NK and T cells.

**Figure 6:**
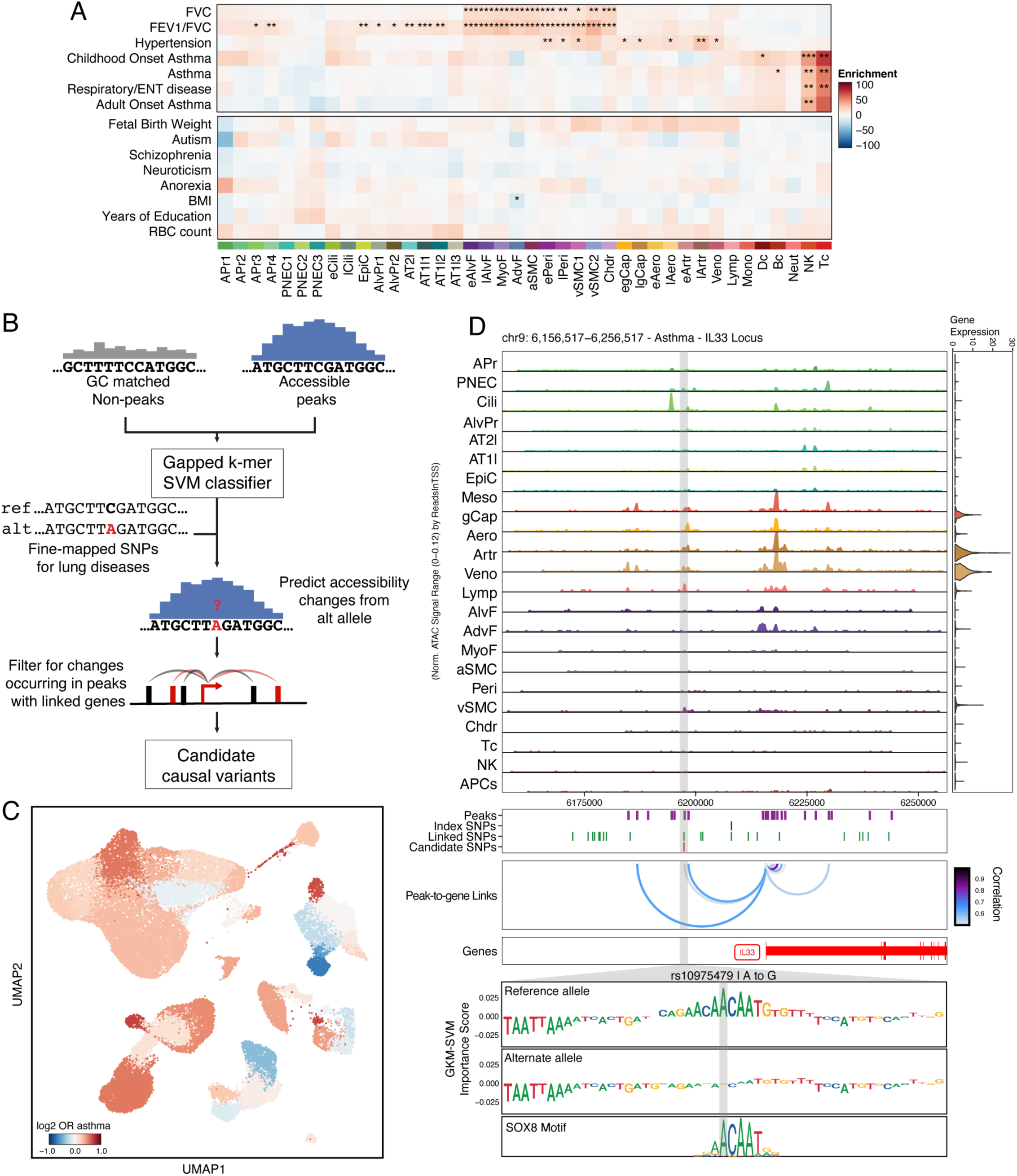
Nominating candidate causal variants of pulmonary diseases. A) Enrichment of GWAS SNPs for pulmonary and non-pulmonary traits and diseases in cell type specific peaks using LDSC. FDR corrected P values from LDSC enrichment tests are overlaid on each cell of the heatmap (*FDR < 0.05, **FDR < 0.005, *** FDR < 0.0005). B) Schematic for candidate causal variant prioritization using gapped k-mer support vector machine (gkm-SVM) classifier. C) Enrichment of GWAS SNPs for asthma for compared to random SNP in peaks specific to each cluster. Each cluster is colored based on odds ratio of enrichment for asthma GWAS SNPs over random SNPs. D) Predicted accessibility change from rs10975479 at the IL33 locus. Pseudobulked chromatin accessibility tracks around IL33 locus are shown with the violin plot for expression of IL33 for each cluster. Accessible peaks, GWAS lead SNP, fine-mapped SNP candidates in LD with lead SNP, and candidate functional variant are shown in the box below the chromatin accessibility tracks. GkmExplain importance scores for 50bp region around rs10975479 for the reference and alternate alleles. The SOX8 motif is the most significant motif matching this change in importance score.

Next, we used the cluster-specific peaks to train a support vector machine (SVM) classifier to nominate functional causal variants. We trained this computationally light- weight cluster-specific gapped k-mer SVM (gkm-SVM) model based on cluster specific accessible peaks versus nucleotide GC content matched non-peaks to predict accessibility from sequence (Figure 6B) (Ghandi et al., 2014, 2016). These models were stable across 10-fold cross validation scheme on held out chromosomes (Supplementary Figure 7). Then, using GkmExplain, we interpreted the impact of each variant allele on disrupting accessibility based on our cluster specific models and filtered for alleles occurring in peaks linked to genes. Using this strategy, we prioritized asthma SNPs that were enriched in immune clusters but also in endothelial clusters (arteries and veins) and stromal clusters (airway smooth muscle cells, adventitial fibroblasts) (Figure 6C). These prioritized SNPs with predicted motif changes and their target genes are available in Supplementary Table 1. One of the prioritized SNPs (rs10975479, A to G variant) was predicted to have a strong disruption of a SOX8 motif in a peak linked to expression of IL-33 (Figure 6D). IL-33 expression was mainly observed in arterial and venous cells, consistent with the understanding that IL-33 functions as an endothelial barrier disruption signal due to tissue injury or infectious agents (Saikumar Jayalatha et al., 2021). Furthermore, recent clinical trials have shown promising results for asthma control and quality of life for IL-33 blockade with itepekimab in asthma patients (Wechsler et al., 2021). Therefore, the cell type specific sequence to accessibility models provide prioritization for likely disease relevant variants of the human lung.

## Discussion

We created a multiomic atlas of human pulmogenesis by simultaneously capturing both the transcriptome and the chromatin accessibility profile from developing human lungs from(PCW 12-23). We then identified gene regulatory networks to characterize novel cell types in lung development, nominate transcription factor (TF) regulators of cell states, describe cell type specific transcription factor usage of enhancers, and prioritized cell types and variants responsible for pulmonary physiology and pathophysiology. By linking highly regulated genes (HRGs) to highly regulating peaks (HRPs) and developing interpretable cell type specific convolutional neural network models, we provide a framework and a rich resource to identify enhancers, their target genes, and associated TF motifs at specific enhancers and cell types across lung development.

Along the differentiation trajectories of aerocytes and arterial endothelial cells, we nominated candidate transcription factor regulators. One of our nominated aerocyte transcription factors, FOXF1, has been shown to be mutated in alveolar capillary dysplasia, therefore suggesting that it is necessary for aerocyte differentiation. We also nominated MEF2A and STAT4, which have been previously implicated in vascular development broadly. Specifically, MEF2A has been shown to be necessary for sprouting angiogenesis, particularly being necessary for the density of projections of retinal vasculature (Sacilotto et al., 2016). Furthermore, the loss of STAT4 has been shown to cause developmental failure of pulmonary arterial vasculature in zebrafish (Meng et al., 2017). For arterial cell types, we recapitulated many of the known SOX transcription factors but also nominated KLF12. While other KLF family members such as KLF2 and KLF4 have been shown to be necessary for arterial endothelium fates (Sangwung et al., 2017; Sweet et al., 2023), KLF12 has mainly been studied in the context of epigenetic dysregulation of carcinogenesis (Godin-Heymann et al., 2016; Li et al., 2023). Given the correlated expression and motif activity in arterial differentiation fate, KLF12 may warrant further study in its role in endothelial differentiation. Our trajectory analyses implicate new transcription factors for their potential role in aerocyte and arterial differentiation, broadening candidates for in vitro differentiation models and disease processes.

We identified novel vascular smooth muscle cell populations and nominated distinct functions, anatomic associations, and reciprocal signaling responsible for their development. A previous report inferred that these cell types were intermixed in all vascular structures (He et al., 2022). However, we show that these cell types are spatially distinct and not intermixed, with vSMC-a surrounding arteries and vSMC-v surrounding veins. These cell types and localizations are distinct at all examined fetal time points. However, in the adult human lung, only a single vascular smooth muscle cell population has been reported (Travaglini et al., 2020). Therefore, the two distinct vSMCs function during development as a secondary response to associated endothelial differentiation or may be a necessary signaling partner to amplify artery vs. vein endothelial differentiation and subsequently coalesce into a common vSMC cell type postnatally. It is also possible that adult pulmonary vSMC subpopulations exist but have not been sufficiently captured or profiled to differentiate the subtypes. Given that pathophysiology of pulmonary hypertension depends on dysfunctional vSMC proliferation and neointimal lesion formation (Saygin et al., 2020; Steffes et al., 2020), further characterization of these niches in the developmental context may reveal mechanisms hijacked in disease.

Along the differentiation trajectory towards surfactant producing AT2 cells, we identified transcription factors with high enrichment of motifs in the CREs linked to surfactant production. We observe the highest enrichment for the glucocorticoid receptor (NR3C1) in these CREs, proposing a direct mechanism for antenatal steroid use to promote lung maturation and surfactant production in premature deliveries (Liggins, 1969; Liggins and Howie, 1972). Furthermore, this strategy nominated additional transcription factors including the SOX transcription factors and NR3C2, the mineralocorticoid receptor, as potential therapeutic targets to further advance lung maturation and surfactant production. Because the time points present within this study are limited to the earliest stages of alveolar maturation, these nominated transcription factors represent the earliest regulators. Further perinatal time points may help to uncover additional regulatory transcription factors.

Finally, we prioritized variants of pulmonary function and asthma and nominated candidate TF motifs disrupted by the variants using sequence to chromatin accessibility machine learning models trained on cell type specific chromatin accessibility profiles. While this strategy provides a powerful framework to link variants to disease phenotypes, additional experimental validation within the appropriate cell type will be necessary to definitively prove the impact of these variants on gene regulation. Furthermore, while congenital heart disease and developmental neurologic conditions such as autism spectrum disorder have significant efforts in cataloguing de novo variants in large cohorts, similar scale efforts in developmental lung disorders such as the risk of bronchopulmonary dysplasia will be necessary. Future large scale de novo variant identifications for developmental lung disorders can leverage the strategy and the atlas we present in this work to nominate and validate mechanisms directly across all cell types of the developing lung.

In summary, our multiomic atlas of human lung development not only uncovers new regulatory mechanisms and cell types but also serves as a foundational reference to inform both developmental biology and the pathogenesis of lung disease.

## Methods

### Human tissue, ethics, and institutional approval

De-identified tissue samples were obtained at Stanford University School of Medicine from elective pregnancy terminations under a protocol and informed consent for the research use of tissues approved by the Research Compliance Office at Stanford University. No demographic information was collected. Consent was obtained by the medical team. The relevant tissue sample processing and analyses were performed under protocol SCRO-796, approved by the Stem Cell Research Oversight Panel (SCRO) at Stanford University. Lung tissue samples at PCW12 to PCW23 were immediately transferred to 1x DPBS on ice following a protocol adapted from previous protocol (Pașca et al., 2019) and processed for single cell analyses or processed for immunofluorescence within 3 h of the procedure.

### Single cell dissociation

Tissue dissociation was performed with modifications from previously described adult lung dissociation (Travaglini et al., 2020). Briefly, ∼200mg of tissue was chopped with sterile scissors on ice and transferred to a gentleMACS type C tube with 400µg/mL Liberase DL (Sigma 5466202001) and 100µg/mL elastase (Worthington LS002294, dissolved in PBS) in 6mL RPMI (Gibco 72400047). Program m_lung_01.01 was performed on the genlteMACS at room temperature. The sample was then incubated at 37°C for 30min with turning. Program m_lung_02.01 was performed on the gentleMACS at room temperature. Enzymatic digestion was neutralized by adding 6mL 5% FBS in PBS. DNAseI digestion was performed by adding 0.33U/mL DNAseI (Worthington LS006343) and incubating at 37°C for 5 min. Sample was centrifuged at 300xg for 5min at 4°C. Cell pellet was resuspended in 5mL 5% FBS in PBS and filtered through a 70µm filter (Miltenyi 130-098-462). Sample was centrifuged at 300xg for 5min at 4°C. Red blood cells were removed using 2mL ACK lysis buffer (Gibco A1049201) to resuspend pellet and incubate on ice for 1 min. ACK lysis was neutralized using 5mL ice cold 5% FBS in PBS. Sample was centrifuged at 400xg for 10min at 4°C and the pellet was resuspended in 1mL ice cold 5% FBS in PBS for counting.

### Multiome data generation

Nuclei preparation was performed with modifications from previous ATAC-seq nuclei preparation protocol (Corces et al., 2017) according to compatible buffers per demonstrated protocol from 10x genomics (10x Genomics CG000365 Rev B). Minimum of 100,000 cells were prepared at a time per sample. Cells were rinsed with 1mL PBS + 0.04% BSA. 100µL chilled lysis buffer (10mM Tris-HCl pH 7.4, 10mM NaCl, 3mM MgCl_2_, 0.1% Tween-20, 0.1% NP-40, 0.01% Digitonin, 1% BSA, 1mM DTT, 1U/µL Protector RNase inhibitor) was used to resuspend cell pellet and incubate on ice for 3 min. Lysis was quenched by adding 1mL chilled wash buffer (10mM Tris-HCl pH 7.4, 10mM NaCl, 3mM MgCl_2_, 0.1% Tween-20, 1% BSA, 1mM DTT, 1U/µL Protector RNase inhibitor). Sample was pelleted at 500xg for 5min at 4°C. Wash was repeated three times. Nuclei was resuspended using diluted nuclei buffer (1x Nuclei Buffer, 1mM DTT, 1U/µL RNase inhibitor) and cell concentration was adjusted to target 9000-10000 nuclei per lane. Library generation was performed according to Chromium Next GEM Single Cell Multiome ATAC + Gene Expression user guide (10x Genomics CG000338 Rev E).

### Multiome data processing

Raw sequencing data were converted to fastq files using “cellranger-arc mkfastq” (10x Genomics, v2.0.0). scRNA-seq and scATAC-seq reads were aligned to respective reference files based on GRCh38 (10x Genomics, GRCh38-2020-A-2.0.0). Feature count matrices and fragment files for scRNA-seq and scATAC-seq, respectively, were prepared using “cellranger-arc count” (10x Genomics, v2.0.0).

#### Gene expression data quality control, dimensionality reduction, and clustering

Each modality of the multiome data was processed independently while synchronizing across modalities for any cell filtering steps to ensure that every cell includes data from both modalities. All subsequent analyses were performed using R v4.0.2 unless otherwise indicated. Gene expression feature count data was further processed using the Seurat R package (v4.3.0) (Hao et al., 2021). Initial quality control was performed on each sample independently. For each sample, cells with less than 200 genes or 1000 unique molecular identifiers (UMIs) or with more than 20% mitochondrial reads were removed. SCtransform (v2) was performed on each sample independently (Choudhary and Satija, 2022). Consensus set of variable features were identified across all the samples while removing genes more likely to be variable due to technical artifact or sex differences (mitochondrial genes, sex chromosome genes, S phase correlated genes, G2M phase correlated genes). This consensus variable feature set of 3000 genes was used to perform SCtransform (v2) on the merged object across all the samples. PCA was performed using “RunPCA” with the consensus variable features. Uniform manifold approximation and projection (UMAP) implementation from Seurat (n.neighbors = 50, mindist = 0.5, distance metric = cosine) was used to create a two-dimensional representation. Nearest neighbors were identified using “FindNeighbors” with 40 PCs. Clusters were identified using “FindClusters” with default settings.

#### Subclustering and cluster annotations

For cluster annotation, each cluster was manually categorized into compartments based on compartment marker gene expression (Epithelial: EPCAM, CDH1; Endothelial: PECAM1, CLDN5; Stromal: COL1A1, COL1A2; Immune: PTPRC). At this stage, if a cluster expressed multiple incompatible compartment marker genes and showed artifactual patterns in UMI counts or number of genes per cell, the cluster was removed. Subsequently, consensus variable feature selection, dimensionality reduction, and clustering was repeated as needed until all low quality and multiplet-like clusters were removed. Additional subclustering was performed by separating out each compartment and performing the above-described preprocessing procedure (consensus variable feature selection, normalization, dimensionality reduction, and clustering). Subclusters with biologically meaningful differences in gene expression were retained based on literature review. Otherwise, the subclusters were collapsed based on parent cluster identity. Final feature counts matrix for gene expression was stored as a Seurat object for further analysis.

#### Chromatin accessibility quality control, dimensionality reduction, and peak calling

ATAC fragments were processed using the ArchR R package (v1.0.3) (Granja et al., 2021). For each sample, cells with less than 1000 fragments or TSS enrichment of less than 5 were removed. Tile matrix and gene score matrices were generated using the functions “addTileMatrix” and “addGeneScoreMatrix”. Finalized gene expression matrix was imported into the ArchR project using “addGeneExpressionMatrix”. Dimensionality reduction was performed using “addIterativeLSI” (varFeatures = 50,000, dimsToUse = 1:30). Clustering was performed with “addClusters” (method = “Seurat”, resolution = 0.6). Low quality clusters or clusters that expressed marker genes that were biologically incompatible were removed. Cluster annotations from processing gene expression data were imported into the ArchR project using the function “addCellColData”. Peak calling was performed using MACS2 with ArchR with the function “addReproduciblePeakSet”. Fine grained cluster annotations performed using gene expression data were used to group the cells for peak calling with maximum number of peaks per cell at 500 (peaksPerCell = 500). The cell by peak matrix was stored in the ArchR project for further analysis.

#### ChromVAR motif analysis

ChromVAR (v1.12.0) (Schep et al., 2017) was used to obtain TF motif accessibility profiles. Position weighted matrices for motif annotations was based on a curated set of non-redundant 1,141 human unique regulators and motifs (Kartha et al., 2021). Using the ArchR function “addMotifAnnotations”, all motif matches in peakset were identified. Then “addBgdPeaks” was used to identify accessibility and GC matched background peaks. Finally, “addDeviationsMatrix” was used to calculate motif deviation z-scores for each of the motifs and this chromVAR motif matrix was stored in the ArchR project for further analysis.

### Data analysis

#### Linking accessible chromatin elements to gene expression

Accessible regulatory regions were linked to candidate target genes (peak-to-gene linkages) by using a correlation based strategy as previously described (Corces et al., 2018). Specific implementation was performed with the ArchR function “addPeak2GeneLinks”. Briefly, up to 500 partially overlapping pseudobulks of up to 100 k-nearest-neighbor single cells for each modality (scRNA and scATAC) were generated based on an approach adopted from Cicero (Pliner et al., 2018). For each pseudobulk, peak counts and gene expression counts were summed. Candidate peak-gene pairs were generated by identifying peaks that are up to 250kb away from the TSS of each gene. Pearson correlation coefficient of log2-normalized accessibility and gene expression counts were calculated for each pair. Pearson correlation coefficient cutoff of 0.5 was used to filter out low quality peak-to-gene links. Additional peak-to-gene links were identified by performing the above-described strategy on each compartment separately. A consensus peak-to-gene link set was created by deduplicating across all peak-to-gene linkages created at different resolutions as previously described (Ober-Reynolds et al., 2023). A final correlation coefficient filter was performed on the final set of non-redundant peak-to-gene linkages.

#### Highly Regulated Gene (HRG) analysis

All expressed genes were ranked based on the number of peak-to-gene linkages. Genes with ≥ 20 linked peaks were defined as highly regulated genes (HRGs). A total of 1,163 HRGs were identified. We performed hypergeometric enrichment test of the HRGs against a previously described set of superenhancer-associated genes (Hnisz et al., 2013). For each of the k-means clusters, we performed Gene Ontology (GO) term enrichment analyses for the top 250 genes with the highest number of linked peaks using topGO (v3.2). Here and in all other subsequent GO term enrichment analyses, enrichment was performed with a foreground set of genes of interest against a background set of all expressed genes. For enrichment of highly regulated peaks (HRPs) in cluster marker peaks, HRPs were first defined as the set of peaks linked to each HRG. Then “peakAnnoEnrichment” was used to perform enrichment of cluster specific marker peaks across HRPs with the expectation that cluster specific accessibility represents cluster specific usage of HRPs and subsequently expression of the linked HRG.

#### Differential cell type abundance testing

Differential cell type abundance testing for clusters across developmental time was performed using the miloR R package (Dann et al., 2022). K-nearest neighbor graph was generated with a minimum of 50 cells and k = 50 for the “buildGraph” function. Then “makeNhoods” was performed with a prop = 0.1. Differential abundance testing was performed using the function “testNhoods”. Neighborhoods were identified as belonging to a given cluster if the neighborhood has greater than or equal to 70% of cells from a single cluster, otherwise the neighborhood was marked as “Mixed”.

#### Differential gene expression

For differential gene expression testing between clusters, DESeq2 (v1.46.0) was used on pseudobulked data (Love et al., 2014). In short, counts from cells were pseudobulked by both clusters and sample of origin. Then differential expression testing was performed using samples as replicates with log fold change shrinkage to correct for changes in genes with high dispersion (Zhu et al., 2019). Log 2 fold change of greater than or equal to 1 and an adjusted p-value threshold of less than or equal to 0.01 was used to identify significantly differential genes between clusters.

#### Motif enrichment analysis

Motif enrichment analysis was performed where appropriate using the ArchR function “peakAnnoEnrichment” with default parameters for FDR ≤ 0.1 and log2 fold change ≥ 0.5.

#### Trajectory analysis

For trajectory analysis, we first calculated the probability of each cell to reach a terminal state using CellRank, a python package (Lange et al., 2022). In short, transition probabilities for each cell was calculated as a weighted mean of the ConnectivityKernel, cell to cell similarity based on transcriptomic similarity, and the RealtimeKernel, cell to cell coupling across developmental time points using optimal transport (Schiebinger et al., 2019b). Then cells were assigned to macrostates using the function “compute_macrostates” with terminal states identified using the function “predict_terminal_states”. Then fate probabilities, which is the likelihood of each cell to transition into each of the terminal macrostates, were calculated using the function “compute_fate_probabilities”. Based on the fate probabilities of each cluster towards the terminal macrostates, trajectory across both RNA and ATAC manifolds were generated using the ArchR function “addTrajectory”. To identify transcription factors likely contributing to development along a trajectory, we correlated transcription factor expression and chromVAR motif deviation scores along the trajectory using the ArchR function “correlateTrajectories”.

#### Ligand receptor analysis

For ligand receptor co-expression analysis, CellPhoneDB database v4.1.0 was used. In short, expressed pairs of ligands and receptors were filtered based on differential expression for at least one of the clusters with a log 2 fold change of greater than 1 and adjusted p-value less than 0.01. Furthermore, clusters were further grouped according to their spatial proximity. We filtered ligand receptor pair expression occurring in cluster pairs that were not observed to be proximal from imaging. Relevant interactions were identified using the function “cpdb_degs_analysis_method” (Garcia-Alonso et al., 2022).

#### ChromBPNet model training and deriving contribution scores

ChromBPNet is a convolutional neural network that uses one-hot-encoded DNA sequence (A = [1,0,0,0], C = [0,1,0,0], G = [0,0,1,0], T = [0,0,0,1]) to predict single base resolution read count profiles. Each cluster-specific model takes in the 2,114 bp region centered around the summit of each pseudobulked ATAC-seq peak and predicts the Tn5 insertion counts at each base pair for the 1,000 bp region centered around the summit. Detailed model architecture is described previously (Ameen et al., 2022; Pampari et al., 2025; Trevino et al., 2021). Fragment files for each pseudobulked cluster was prepared using the ArchR function “getGroupFragments”. Clusters with less than 5 million fragments were removed from further model training. Peak sets for each cluster was prepared by identifying which of the peaks were originally called for a given cluster prior to iterative consensus cluster generation per ArchR. These peaks were filtered for overlap with blacklisted regions extended by 1057 bp on both sides. Chrombpnet pipeline v0.1.7 was used for further model training and predictions. Set of non-peak regions that are GC-matched to the peak set using the “chrombpnet prep nonpeaks” command with default parameters. A 5-fold chromosome hold-out cross-validation scheme for training, validation, and testing was used. Test chromosomes: fold 0: [chr1, chr3, chr6], fold 1: [chr2, chr8, chr9, chr16], fold 2: [chr4, chr11, chr12, chr15, chrY], fold 3: [chr5, chr10, chr14, chr18, chr20, chr22], fold 4: [chr7, chr13, chr17, chr19, chr21, chrX]. Validation chromosomes: fold 0: [chr8, chr20], fold 1: [chr12, chr17], fold 2: [chr22, chr7], fold 3: [chr6, chr21], fold 4: [chr10, chr18]. The Tn5 bias model was trained using the command “chrombpnet bias pipeline” based on the egCap cluster with a bias hyperparameter of 0.5. Bias model performance was evaluated for appropriate Tn5 bias motif learning in both counts and profiles. Cluster specific ChromBPNet models were trained using the command “chrombpnet pipeline”.

To evaluate model performance for counts predictions, we calculated the Pearson correlation coefficient between the predicted and observed log counts for all peak regions. To evaluate model performance for profile shape predictions, we calculated the Jensen Shannon Distance between observed and predicted base-resolution probability profiles for each peak region, with lower divergence being better performance.

To derive cluster specific predicted accessibility contribution at a single base pair resolution, DeepLIFT algorithm was applied (Shrikumar et al., 2017). By interrogating each ChromBPNet models to understand the features learned, we can estimate the contribution of each base pair to the predicted accessibility for that given cell type. Ultimately, this strategy enables a cell type specific base pair resolution contribution score for accessibility. Contribution scores were derived using the command “chrombpnet contribs_bw”. These contribution scores were then averaged across the 5- folds of cross-validation.

#### LD Score Regression

LDSC (v.1.0.1) was used to estimate heritability of pulmonary traits from each cluster. As input annotation for LDSC, cluster specific peaks were generated by identifying which peaks were originally called from a given cluster by overlapping union peak set with the MACS2 peak calls from specific clusters. Peaks that were not identified in more than 25% of all clusters were retained to filter out common peaks across majority of cell types. Summary statistics for partitioning was downloaded from https://console.cloud.google.com/storage/browser/broad-alkesgroup-public-requester-pays/sumstats_formatted. Celltype specific partitioned heritability analysis was performed using 1000 genome EUR phase 3 population reference and the hg38 baseline model (v.2.2). The “ldsc.py” script was used to calculate partitioned heritability in cluster specific peak sets for each pulmonary traits. Benjamini-Hochberg FDR correctio nwas used to adjust heritability enrichment p-values.

#### gkm-SVM training and testing

Previously published strategy for training gapped k-mer support vector machine (gkm-SVM) models using scATAC-seq peaks was used (Corces et al., 2020; Ober-Reynolds et al., 2023). For each cluster, we trained a gkm-SVM classifier to predict whether a genomic sequence is likely accessible or inaccessible in that cell type. First, marker peaks were identified for each cluster using ArchR function “getMarkers” with an FDR cutoff of less than 0.1 and a Log2FC cutoff of greater than 0.5. For training, these peaks were extended to 1001bp regions around summits. Then cluster specific peaks were identified as described above. For negative training data, 4 million random genomic regions of 1001 bp length were identified while masking for assembly gaps and intra-contig ambiguities. For each cluster, negative regions were identified that were matched for GC-content percentile as the marker peaks. A 10 fold chromosome hold out cross validation strategy was used to train and test model performance. Following folds were used for testing: Fold 1: [chr 1], Fold 2: [chr2, chr19], Fold 3: [chr3, chr20], Fold 4: [chr6, chr13, chr22], Fold 5: [chr5, chr16], Fold 6: [chr 4, chr15, chr21], Fold 7: [chr7, chr14, chr18], Fold 8: [chr11, chr17, chrX], Fold 9 [chr9, chr12], Fold 10: [chr8, chr10]. For each fold, chromosomes not used for testing was used for training. SL-GKM package was used with the following options for the “gkmtrain” function (Ghandi et al., 2016; Lee, 2016).The wgkmrbf kernel (t = 5), a word length of 11 (l= 11), 7 informative columns (k = 7), up to 3 mismatches to consider (d = 3), an initial value of 50 for the exponential decay function (M = 50), a half-life parameter of 50 (H = 50), and a precision parameter of 0.001 (e = 0.001). Performance of trained models were assessed from cross-validation folds by calculating AUROC and AUPRC using the ‘PRROC’ R package (v1.3.1) with negative testing data downsampled to match the number of positive testing data sequences.

#### Candidate causal variant nomination

Fine-mapped SNPs from GWAS for asthma (adult onset and child onset from PICS2) and randomly selected fine-mapped SNPs from any GWAS (PICS or Finucane data) were collected. SNPs were then filtered based on overlap with any peak region. Then SNPs were filtered based on fine-mapping posterior probability of ≥ 0.01. This resulted in 1027 SNPs for asthma across both adult and childhood onset. 2500 random SNPs were identified as negative SNP set. For each SNP, 250bp surrounding region was identified and a synthetic alternative allele sequence was created by replacing the reference allele at the middle of the region. Then celltype specific gkm-SVM models were used to calculate GkmExplain importance scores for each base of the reference and alternate allele sequences (Shrikumar et al., 2019). The delta score for each SNP and cell type was calculated by summing the importance scores for the central 50bp of both the reference and alternate allele sequences then subtracting the alternate allele score from the reference allele score.

Statistical significance testing and prioritization was performed for each candidate SNP. First, 3 di-nucleotide shuffled sequences were generated using the “fasta-dinucleotide-shuffle.py” script from the MEME suite (v5.4.1). A reference and alternate allele sequences were generated by replacing the central position of the shuffled sequences with each SNP reference or alternate allele. Then GkmExplain importance scores for each of these shuffled sequences are calculated across all clusters as described above and a delta score is calculated as described above. The delta scores from these shuffled sequences were used as the null distribution for each cluster. The “fitdistrplus” R package (v1.1.6) was used to fit a t-distribution to the delta score null distribution of each cluster and a p-value for each fine-mapped SNP was calculated for each cluster. Next, prominence score was calculated as an estimate of the extent of disruption for transcription factor binding. First, SNPs with a positive delta score were identified, which would suggest that the reference allele is more accessible than the alternate allele. Then, subsequence was identified surrounding each SNP based on importance score in this region exceeding the 97.5^th^ percentile of the di- nucleotide shuffled background. The subsequence boundaries were determined by the position where two consecutive bases had importance scores falling below the threshold. Prominence score was calculated by taking the sum of non-negative GkmExplain importance scores from the reference allele subsequence and then divided by the sum of the non-negative importance scores for the entire surrounding 250bp sequence. Then an exponential distribution was fitted to the null distribution of prominence scores for each cluster and these were used to calculate the prominence p-values for each SNP for each cluster. Together, SNPs were prioritized based on a delta score p-value < 0.05 and a prominence score p-value < 0.05. Then SNPs were filtered again based on whether they fall in a peak that is linked to a gene.

### RNA Fluorescence In Situ Hybridization

To visualize gene expression in situ, RNA fluorescence in situ hybridization (FISH) was performed using the RNAscope protocol (Advanced Cell Diagnostics, Rev B) coupled with post-hybridization antibody staining. After inflation, lungs were fixed in 4% paraformaldehyde at 4°C overnight, then cryo-embedded into Optimal Cutting Temperature, frozen at -80°C and 20um sections were cut using a cryostat. Standard sample preparation and hybridization was performed per protocol. Key changes were made as follows. Sections were post-fixed onto the slide with 10% normal buffered formalin for 15 minutes at room temperature. Protease plus digestion of the sections was performed for 15 minutes. Following completion of ACD protocol, sections were washed twice with PBS + 0.1% Tween-20, blocked for 30 min in preblock consisting of 0.3% Triton X-100, 5% goat serum and 1.5% BSA, and then incubated overnight in primary antibody solution (mouse anti SMA-FITC (1:200; Sigma, F3777) and DAPI (1:1000; Invitrogen, D1306)) at room temperature. The next day slides were washed a mounted with ProLong Gold Antifade (Invitrogen, P36930). Confocal images were captured on a Zeiss 880 Examiner confocal microscope and maximum intensity projections were made and minimally processed using Zen Black software (Carl Zeiss AG).

## Acknowledgements

We acknowledge the members of the Greenleaf Lab, Kuo Lab, Krasnow Lab, and Kumar lab for insightful discussions and advice. We thank members of the Kumar and Kuo labs with support for the tissue sectioning and imaging. S.H.K. was supported by the Paul and Daisy Soros Fellowship. W.J.G. acknowledges support from the Arc Institute and the Chan-Zuckerberg Biohub. M.A.K. is a Howard Hughes Medical Institute Investigator. C.S.K. is a Tashia and John Morgridge Endowed Faculty Scholar of the Maternal and Child Health Research Institute (MCHRI)

## Author Contributions

S.H.K. and W.J.G. conceived of the study. S.H.K., S.H., L.L., A.M.P. designed and performed tissue collection. S.H.K. and S.H. performed sample processing and experiments. S.H.K., S.H., L.S., M.K., and C.S.K. performed tissue sectioning and imaging. S.H.K. processed the data and performed analysis. C.S.K., M.A.K., and WJG supervised the work. S.H.K. and W.J.G. wrote the manuscript, with input from all authors.

## Data availability

All data presented in this manuscript (fragment files, counts matrices, data objects, ChromBPNet models and contribution scores) are deposited at Zenodo and available at https://zenodo.org/communities/hlda/records.

## Code availability

Code for analysis is available at https://github.com/GreenleafLab/HLDA.

---

**Supplementary Figure 1:**
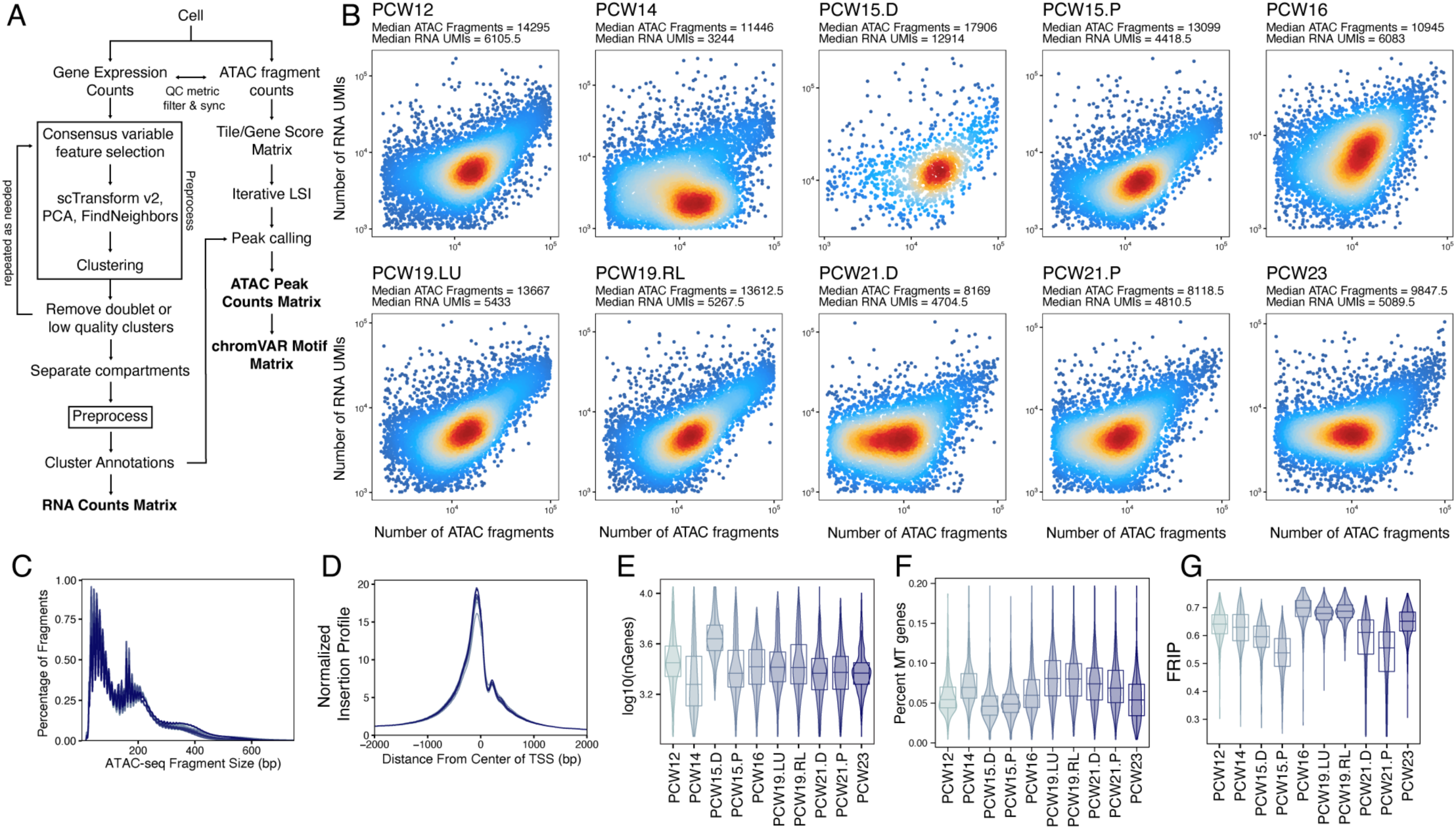
Single cell multiome preprocessing and quality control metrics. A) Schematic for preprocessing pipeline for single cell multiome dataset. B) Sensitivity of the multiomic assay for each sample. Each cell is plotted based on number of ATAC fragments and number of RNA unique molecular identifiers detected. C) Fragment size distribution for each sample. D) Transcription start site enrichment for each sample. E, F, G) Distribution of number of genes (E), percentage of mitochondrial (MT) genes (F), Fragments in Peaks (FRiP) (G) detected per cell for each sample.

**Supplementary Figure 2:**
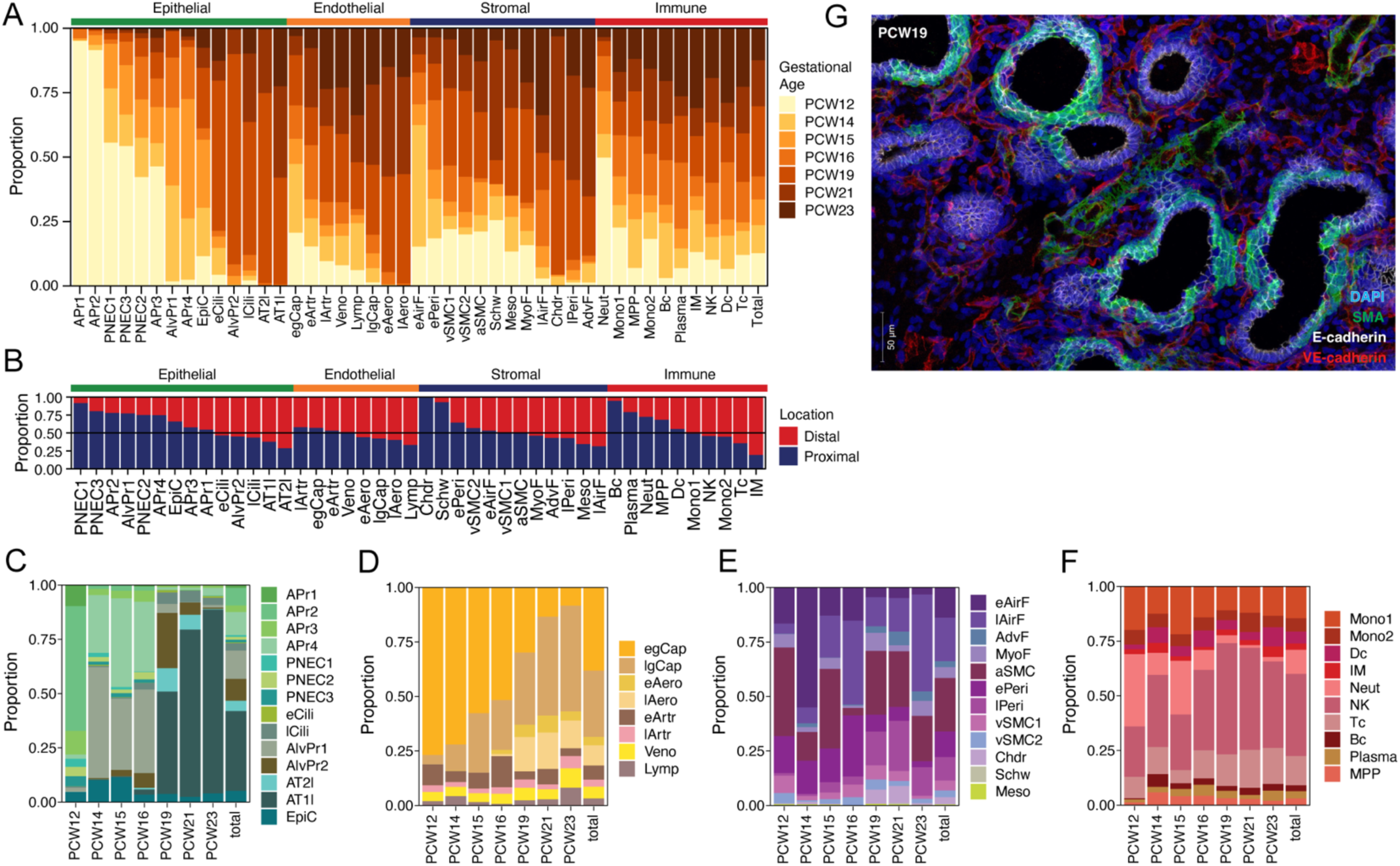
Cluster composition across gestational age, proximal-distal axis, and compartment. A) Distribution of gestational age for each cluster grouped by each compartment and ordered by increasing mean age. B) Distribution of cells from proximal or distal lung when location information is available. C-F) Distribution of clusters for each gestational age for epithelial (C), endothelial (D), stromal (E), and immune (F) compartments. G) Representative immunofluorescence image of PCW19. Pseudocolored blue for DAPI, green for smooth muscle actin, white for E-cadherin (epithelial), red for VE-cadherin (endothelial)

**Supplementary Figure 3:**
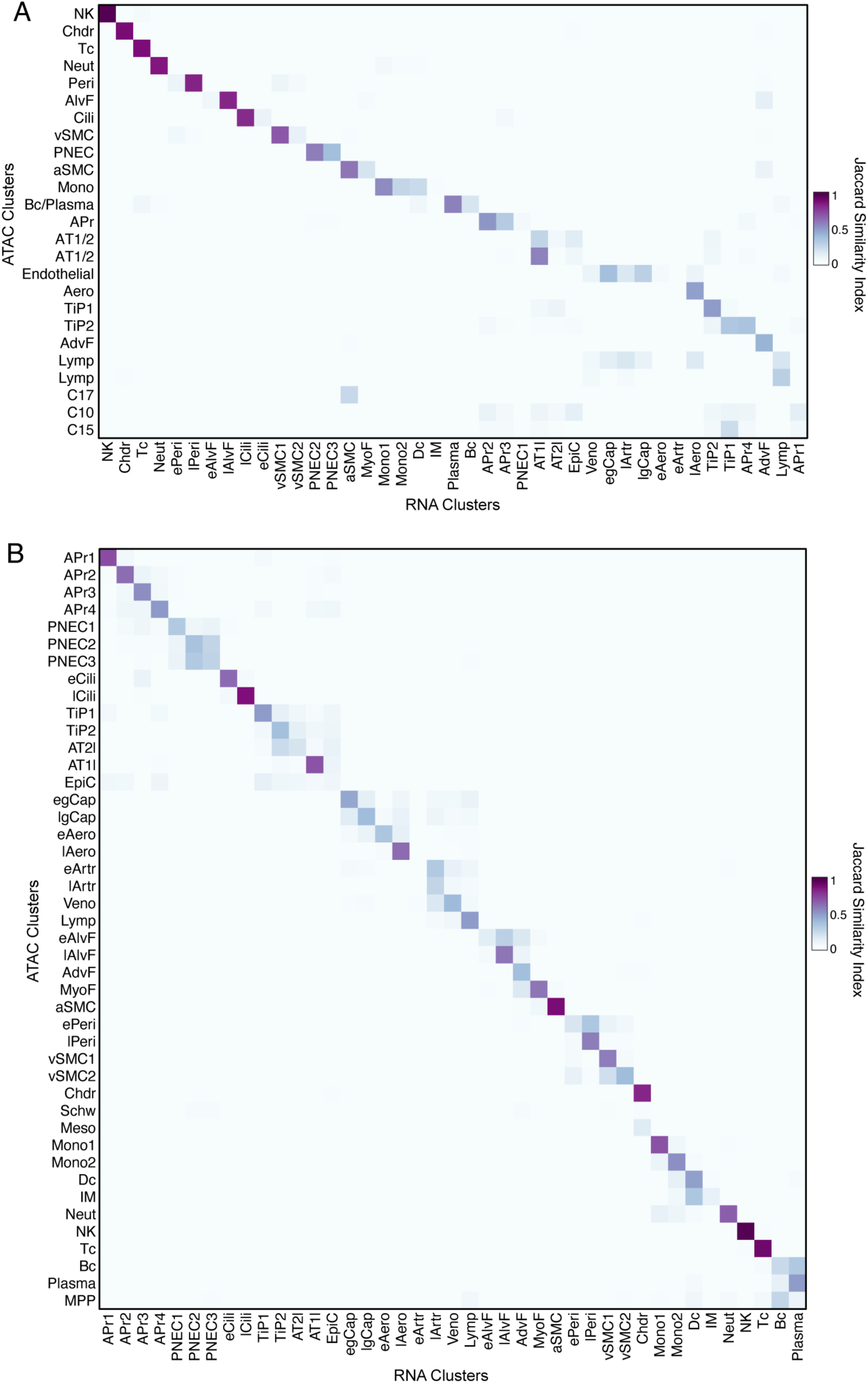
Comparison between true multiome vs. simulated integration in accuracy of cluster identity transfer. A) Correspondence of cluster identity assignments from simulated integration. Heatmap of Jaccard similarity index between cluster identity assignments from simulated integration. B) Heatmap of Jaccard similarity index between cluster identity assignments from direct use of multiome information to transfer cluster assignment based on cell barcode.

**Supplementary Figure 4:**
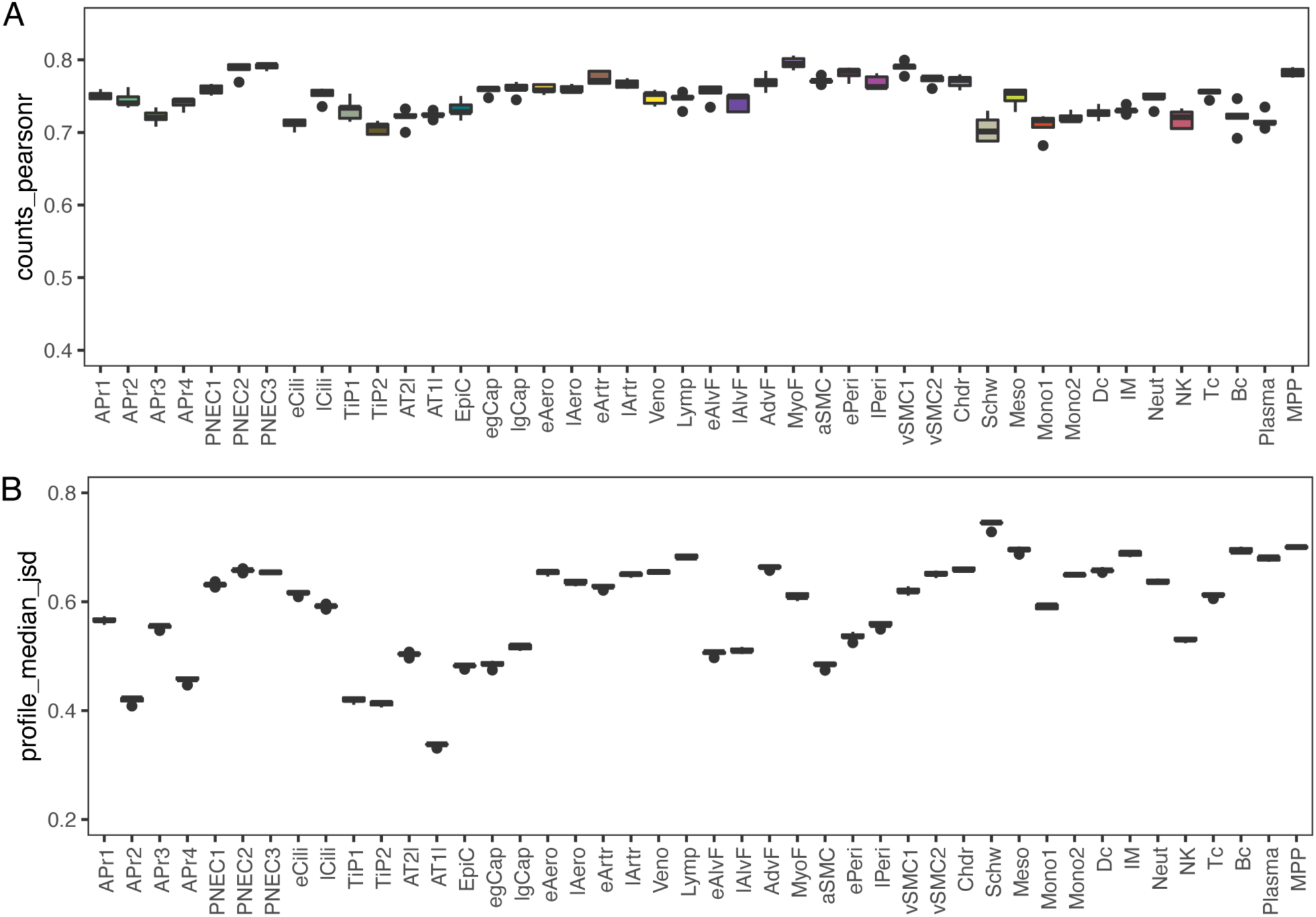
Performance evaluation of cluster specific ChromBPNet models. A) Spearman correlation between observed and predicted total counts in peaks for each model across 5-fold cross-validation. B) Median Jenson-Shannon distance between observed and predicted base resolution profiles at peaks across 5-fold cross-validation.

**Supplementary Figure 5:**
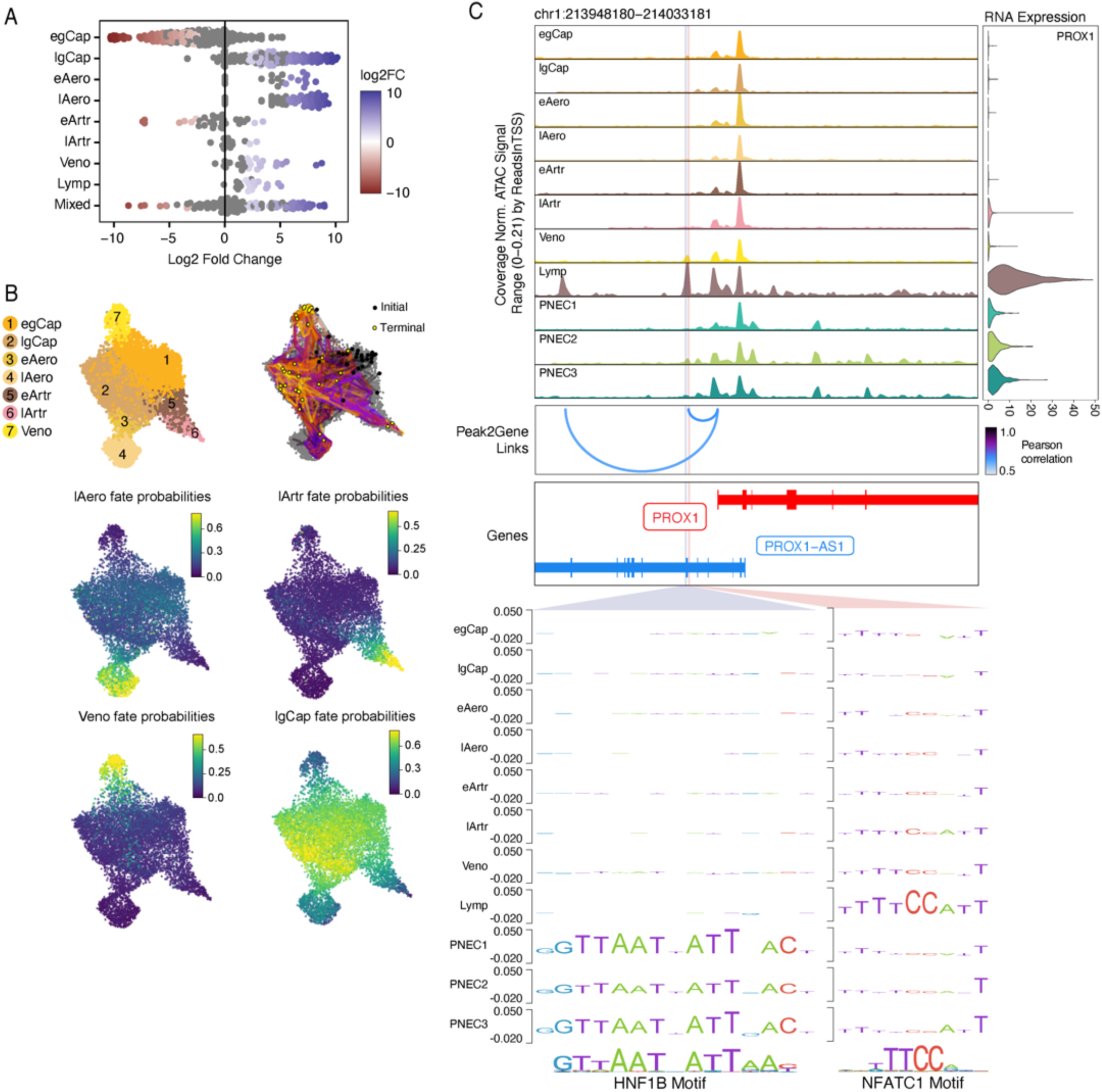
Endothelial trajectory analysis and regulatory dissection of PROX1 enhancer. A) Differential abundance testing of neighborhoods across post conception weeks for clusters in the endothelial compartment B) Fate probabilities for the 4 terminal macrostates within the endothelial compartment. C) ChromBPNet predicted motif accessibility in a PROX1 enhancer validated in mice by Kazenwadel et al. (2023). NFATC1 motif and HNF1B motif accessibility is shown within the same peak linked to expression of PROX1.

**Supplementary Figure 6:**
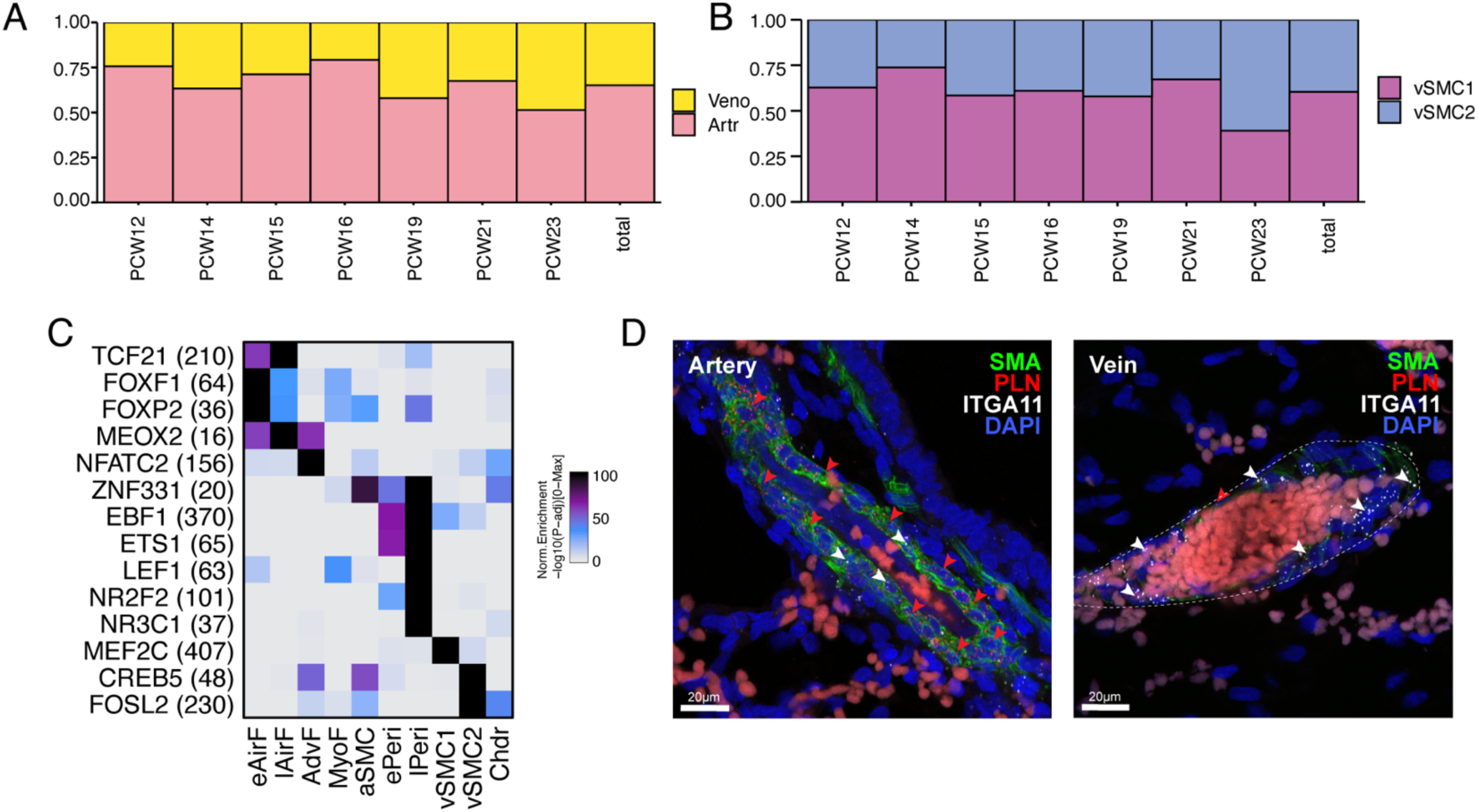
A) Ratio of all arterial compared to venous endothelial cells across development. B) Ratio of vSMC1 and vSMC2 cells across development. C) Heatmap of motif enrichment of expressed transcription factors in marker peaks only within the stromal clusters. D) Representative IF-FISH images of PCW 19 artery and vein with surrounding vascular smooth muscle cells. Pseudocolored with smooth muscle actin antibody in green, FISH probes for PLN in red, ITGA11 in white, and DAPI in blue. Arrows highlight examples of puncta for either PLN (red) or ITGA11 (white).

**Supplementary Figure 7:**
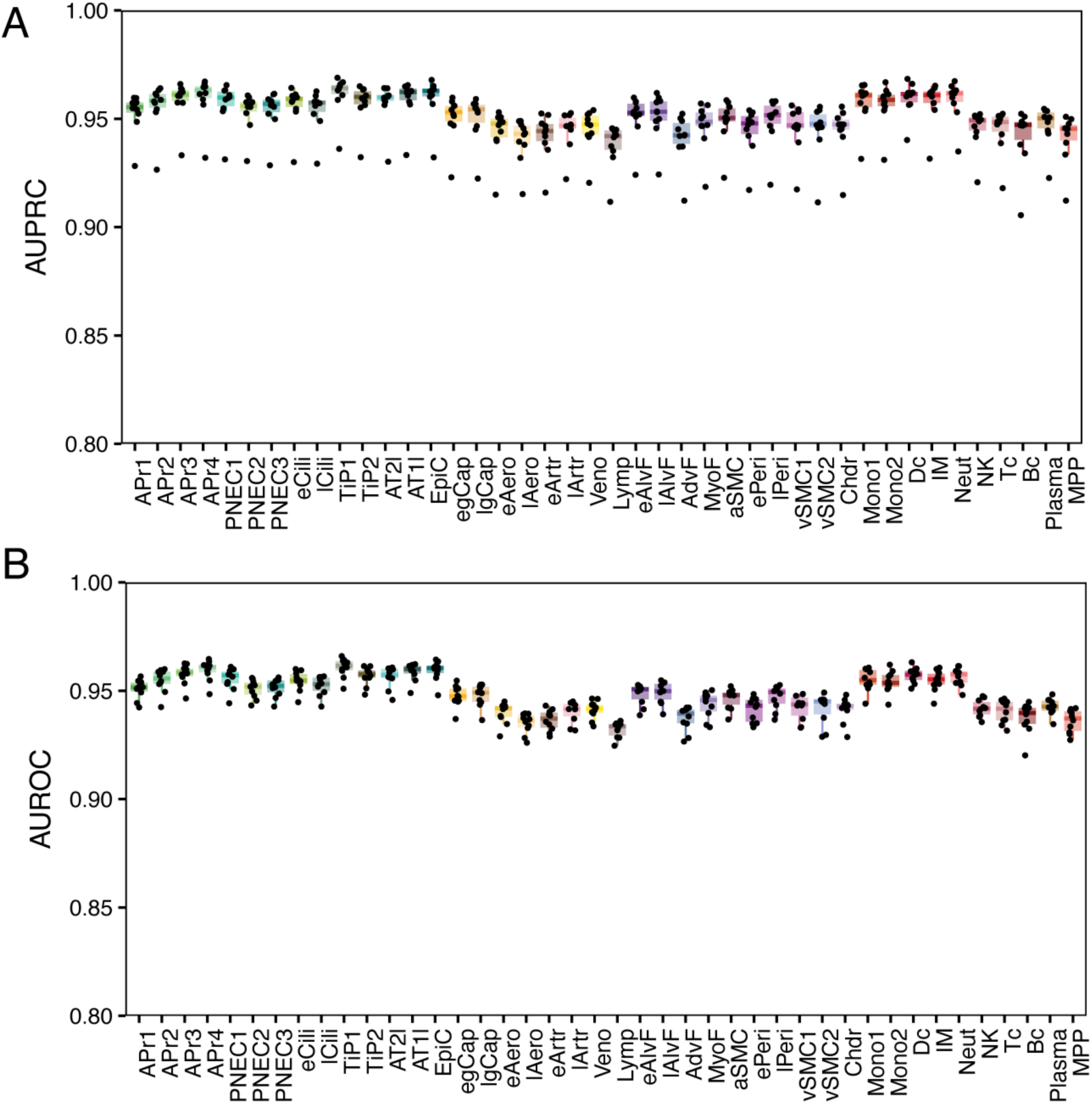
Performance evaluation of cluster specific Gapped k-mer SVM (gkm-SVM) models. A) Boxplot of area under the receiver operator (AUROC) or B) precision recall (AUPRC) for cluster specific gkm-SVM classifiers.

